# Variant-specific spike conformational dynamics shape memory B cell selection during recall

**DOI:** 10.64898/2026.07.18.739204

**Authors:** Manon Broutin, Aurélien Sokal, Alice Dejoux, Delphine Planas, Matthias Vanderkerken, Ignacio Fernández, Imane Azzaoui, Alexis Vandenberghe, Manon Charlet, Atousa Arbabian, Jerôme Megret, Lamine Touré, Cyril Planchais, Laurence Berard, Tabassome Simon, Jean-Claude Weill, Hugo Mouquet, Olivier Schwartz, Pierre Bruhns, Felix A. Rey, Odile Launay, Matthieu Mahévas, Pascal Chappert

## Abstract

Immune imprinting profoundly shapes antibody responses to breakthrough infections and vaccination against evolving endemic viruses. Current vaccine design primarily focuses on antigen sequence and variant mutations but rarely consider the structural context in which these antigens are being recognized. In this study, we make use of the longitudinal analysis of memory B cell (MBC) responses in boosted individuals enrolled in the COVIBOOST clinical trial to provide a proof of principle that variant-specific conformational dynamics can impact MBC recruitment and protective antibody responses, independently of epitope conservation. Combining functional characterization of MBC-derived monoclonal antibodies, repertoire analysis, epitope mapping and *in silico* structural modeling of epitope accessibility, we found that the adjuvanted B.1.351 spike vaccine preferentially recalled MBCs targeting exposed receptor-binding domain neutralizing epitopes. This preferential recruitment arises from the more restricted conformational dynamics of the B.1.351 spike compared to the ancestral Hu-1 spike, leading to increased masking of class 4 and 5 cryptic RBD epitopes. These findings demonstrate that antigen conformational dynamics can be leveraged to redirect pre-existing immunity toward neutralizing epitopes upon boost immunization.

## INTRODUCTION

The rapid development of COVID-19 mRNA vaccines has represented a major turn in mitigating the impact of the SARS-CoV-2 pandemic, lowering disease severity, hospitalizations, and preventing more than 20 million deaths(Ioannidis et al. 2025). Early in the pandemic, the spike protein was identified as the main target of neutralizing B cell responses(Tong et al. 2021; Piccoli et al. 2020). All currently available vaccines in Western countries, including widely used mRNA platforms, initially encoded the ancestral Hu-1 spike protein, which has been periodically updated to reflect circulating SARS-CoV-2 variants(Uraki et al. 2026). Evidence from longitudinal studies have converged on the fact that heterologous spike-based boosts primarily trigger a cross-reactive B cell response. This has been well illustrated following both vaccination(Alsoussi et al. 2023) or infection with Omicron BA.1(Kaku et al. 2023; Sokal et al. 2023) and later variants such as XBB.1.5(Tortorici et al. 2024), where pre-existing memory B cells (MBCs) targeting conserved epitopes are predominantly recruited in the recall response, with no or very little *de novo* naive B cell response against variant specific neo-epitopes so far. This phenomenon, coined immune imprinting, has been described against influenza, where exposure during early life has a life-long impact on subsequent immune responses.

The receptor-binding domain (RBD) of the spike trimeric glycoprotein (S) is the main target of the neutralizing response, accounting for more than 80 % of serum neutralizing antibodies(Piccoli et al. 2020; Tong et al. 2021). The RBD is one of the most mutation-dense and evolutionarily dynamic regions, shaped by intense immune selection and functional constraints(Uraki et al. 2026). Antibody responses against the RBD have been extensively characterized and classified according to the binding pattern of antibodies(Barnes et al. 2020). This classification currently distinguishes class 1, 2 and 3 antibodies - mostly targeting exposed RBD epitopes and frequently exhibit high neutralizing potency - from class 4, 5 and some class 3 antibodies targeting cryptic or less accessible epitopes buried within the spike trimer(Barnes et al. 2020; Fan et al. 2025; Jensen et al. 2023; Cui et al. 2024). Antibodies targeting cryptic epitopes are less frequently neutralizing. However, because they bind to conserved regions of the sarbecovirus spike, they often display broader cross-reactivity(Starr et al. 2021; Jette et al. 2021; Burnett et al. 2021; Jensen et al. 2023; Cui et al. 2024).

Within the canonical prefusion spike trimer, the three RBDs undergo continuous dynamic transitions between receptor-inaccessible “down” and receptor accessible “up” states, with conformational states displaying all RBD in the down position (closed) or 1 or 2 RBDs up (partially open) having been most frequently observed(Benton et al. 2020; Wrapp et al. 2020). The canonical conformational state with all three RBDs in the up position (fully-open) is rare and predominantly captured in ligand-stabilized structures, such as in complex with angiotensin-converting enzyme 2 (ACE2)(Benton et al. 2020). Beyond these canonical states, hydrogen deuterium exchange mass spectrometry (HDX-MS) analyses further revealed distinct expanded-open trimer states characterized by increased exposure of inter-protomer interfaces with various level of loosely-associated ectodomain protomers(Costello et al. 2022; Stuible et al. 2023). The dynamic equilibrium between these various above described states is modulated by variant-specific mutations, such as the increased prevalence of two-RBDs-up conformations in the recombinant Beta spike (B.1.351)(Bruch et al. 2024), and environmental factors, including temperature and pH(Costello et al. 2022; Stuible et al. 2023; Edwards et al. 2021; T. Zhou et al. 2020).

The rapid and highly efficient development of the first SARS-CoV-2 mRNA vaccines was only made possible by extensive prior research on the structure of the coronavirus spike and other class I fusion proteins. Previous studies on MERS-CoV demonstrated that introducing two proline substitutions (S-2P) stabilized the spike trimer in its prefusion conformation and enhanced its immunogenicity(Sanders and Moore 2021). This two-proline stabilization, at residues 986 and 987, has been preferentially used in most first-generation COVID-19 vaccines. Yet, the dynamic equilibrium between the various spike trimer conformational states under physiological conditions, and its impact on anti-RBD antibodies accessibility and underlying B cell recruitment remains poorly defined.

To date, few randomized head-to-head trials comparing vaccines expressing different SARS-CoV-2 spikes have been conducted. One of these phase 3 trials, COVIBOOST(Launay et al. 2022), found that a recombinant vaccine based on the B.1.351 spike elicited higher neutralizing antibody titers against the B.1.351 SARS-CoV-2, but also against both the ancestral and the more distant Omicron BA.1 strains of SARS-CoV-2, than the ancestral spike itself(Launay et al. 2022). This vaccine has further demonstrated clinical effectiveness comparable to that of the bivalent mRNA booster encoding BA.4/5 and ancestral spike against Omicron XBB.1.5(Kirsebom et al. 2024). Such an expansion of immune breadth and anamnestic response has not been observed in other clinical trials comparing vaccines based on different spike proteins, including studies of Omicron BA.1 and BA.5 formulations delivered as either mRNA or recombinant protein vaccines(Bennett et al. 2024; Chalkias et al. 2022). The well-defined antigenic and conformational features of the B.1.351 spike(Bruch et al. 2024) provided a unique framework to determine how these properties reshape an already mature MBC repertoire.

Making use of longitudinal samples from the COVIBOOST trial, we show that the MVB.1.351 vaccine preferentially recruits cross-reactive MBCs targeting neutralizing exposed RBD epitopes. This targeted recruitment is driven by the conformational dynamics of the B.1.351 spike trimer, which less frequently adopts the expanded open trimer state than the Hu-1 spike, limiting access to cryptic RBD epitopes. Our results highlight conformational stabilization of the spike trimer into less dynamic states as a powerful approach to overcome immune imprinting and pave the way for next-generation vaccine strategies.

## RESULTS

### The MVB.1.351 vaccine recruits a specific population of RBD-specific memory B cells

The COVIBOOST clinical trial was a phase III controlled randomized trial (COVIBOOST; NCT05124171; EudraCT number, 2021-004550-33) designed to address the immunogenicity of the preS dTM-AS03 adjuvanted recombinant Beta (B.1.351) spike (MVB.1.351, Sanofi-Pasteur/GSK)(Launay et al. 2022). This variant-targeted vaccine was compared to a preS dTM-AS03 adjuvanted recombinant ancestral Hu-1 spike vaccine (MVD614, Sanofi-Pasteur/GSK) as well as an ancestral Hu-1 spike encoding mRNA vaccine (BNT162b2, COMIRNATY^®^, Pfizer-BioNTech). All three vaccine candidates were administered as a boost in previously immunized individuals with two doses of the BNT162b2 vaccine (see **Methods**). The Hu-1 and the Beta B.1.351 spikes used in both preS dTM-AS03 adjuvanted recombinant protein vaccines differ by only 11 mutations, including 3 within the RBD (K417N, E484K, and N501Y) (**Table 1**). They shared the same engineered two-proline stabilizing mutations at residues 986 and 987, also present in the BNT162b2 encoded Hu-1 spike, as well as additional mutations at the S1/S2 junction (682-685, RRAR to GSAS) to abrogate furin cleavage. Serological follow-up revealed an increased strength and breadth of the response induced by the MVB.1.351 vaccine(Launay et al. 2022). As part of an ancillary study to this trial, aimed at understanding the cellular determinants of this increased humoral response, we analyzed here the underlying memory B cell response in a subset of 46 patients, randomly selected from each arm of the study (**Fig. 1a, Methods,** and **Table S1**). Peripheral blood mononuclear cell (PBMC) samples were available for all patients at 28 days (D28) and/or 3 months (M3) post-boost, and for 7 patients at day 15. The included individuals displayed serological responses representative of the overall trial outcomes (**Fig. 1b**, **Table S1b**, and(Launay et al. 2022)), confirming an efficiently balanced allocation of samples across randomization arms. Notably, those vaccinated with the MVB.1.351 vaccine elicited higher serum neutralizing responses against the Beta (B.1.351) strain at D15 and D28, as compared to individuals vaccinated with the BNT162b2 and the MVD614 vaccines (**Fig. 1b**).

**Fig. 1.**
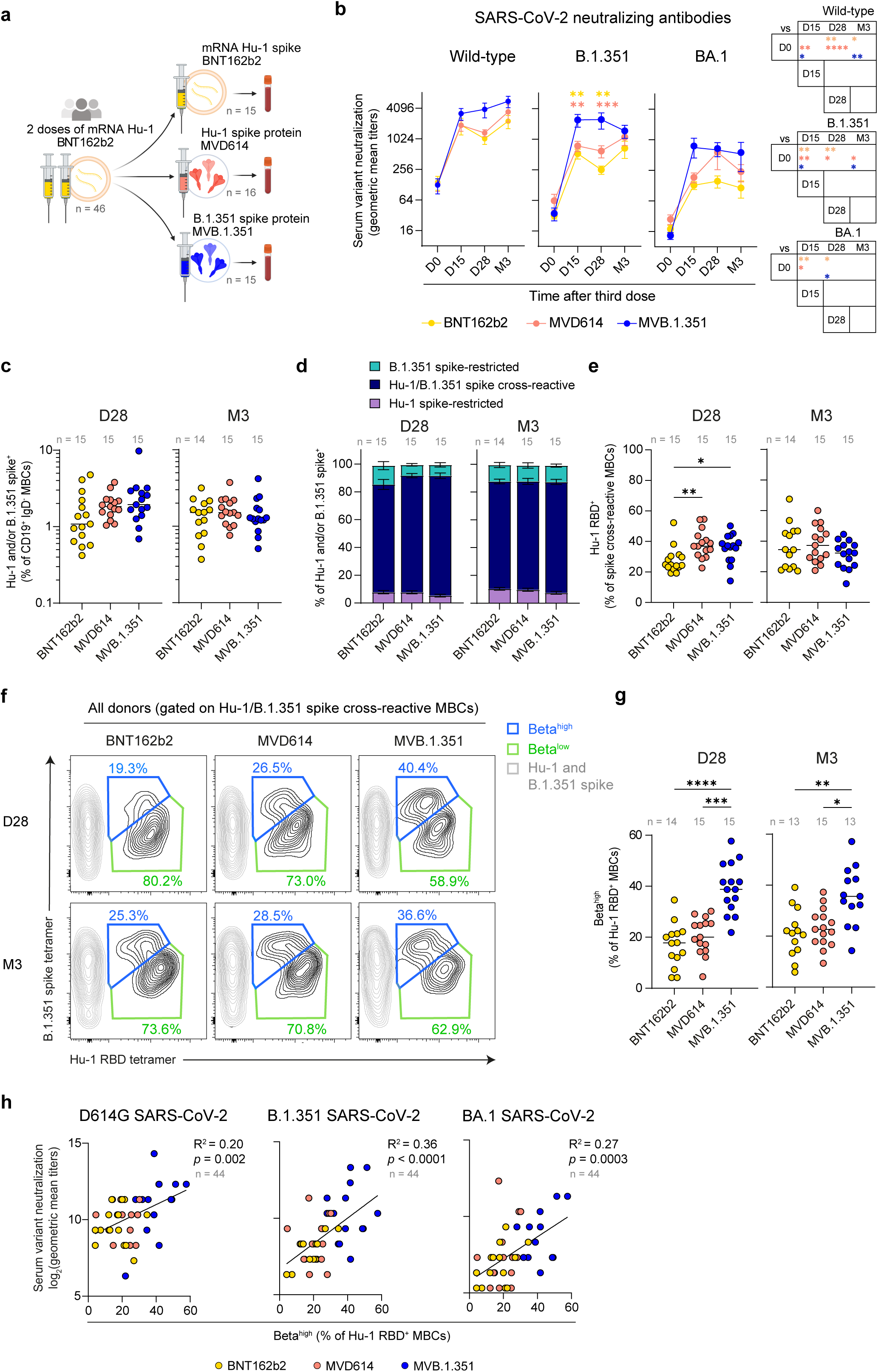
**The MVB.1.351 booster preferentially recruited an RBD-specific memory B cell subpopulation.**(**a**) Overview of the COVIBOOST ancillary study. (**b**) Evolution of geometric mean titers of serum neutralizing antibodies against wild-type (left), B.1.351 (middle), and BA.1 (right) SARS-CoV-2 measured by *in vitro* microneutralization assay after booster vaccination in each group. Data are presented as mean ± SEM. *p*-values displayed within the plots correspond to comparisons between the MVB.1.351 vaccine and other vaccines at each time point, while *p*-values on the right indicate within-group comparisons across time points. (**c**) Frequency of cells recognizing any of the Hu-1 and the B.1.351 spike (Hu-1 and/or B.1.351) among CD19^+^IgD^-^ memory B cells (MBCs) in each donor across groups at D28 and M3 post-boost; bars indicate the median. (**d**) Distribution of cells recognized only the Hu-1 spike (Hu-1 spike-restricted, pastel purple), both Hu-1 and B.1.351 spikes (Hu-1/B.1.351 cross-reactive, dark blue), or only the B.1.351 spike (B.1.351 spike-restricted, cyan) MBCs among total spike-specific MBCs in each group at D28 and M3. Means are shown with SEM. (**e**) Proportion of Hu-1 RBD-specific cells among Hu-1/B.1.351 spikes cross-reactive CD19^+^IgD^-^ MBCs in each donor from each group at D28 and M3 post-boost; bars indicate the median. (**f**) Flow cytometry plots of concatenated donor samples by vaccine group and timepoint depicting Hu-1 RBD and B.1.351 spike tetramer stainings in CD19^+^IgD^-^ Hu-1/B.1.351 spike cross-reactive MBCs. Within Hu-1 RBD^+^ cells, the gating strategies for the Beta^high^ (blue) and the Beta^low^ (green) populations are shown. Values indicate the respective percentage of Hu-1 RBD-specific MBCs in Beta^high^ or Beta^low^ populations. Non-RBD specific cells are recolored in light grey for visualization purpose only. (**g**) Proportions of Beta^high^ CD19^+^IgD^-^ MBCs among Hu-1 RBD- specific MBCs in each donor from each group at D28 and M3 post-boost; bars indicate the median. (**h**) Correlation between log_2_-normalized *in vitro* serum neutralizing geometric mean titers against authentic D614G, B.1.351 and BA.1 SARS-CoV-2 and the frequency of Beta^high^ MBCs among RBD-specific CD19^+^IgD^-^B cells at D28 post-boost in subjects. Dot color indicates vaccine group. (b) Mixed-effects model with Tukey’s correction for comparisons across and within time points. (c and e) Kruskal-Wallis test with Dunn’s multiple-comparison correction. (h) Simple linear regression. \*\*\*\**p* < 0.0001, \*\*\**p* < 0.001, \*\**p* < 0.01, \**p* < 0.05. See also **Fig. S1**, **Tables S1 and S2**.

**Table 1:**
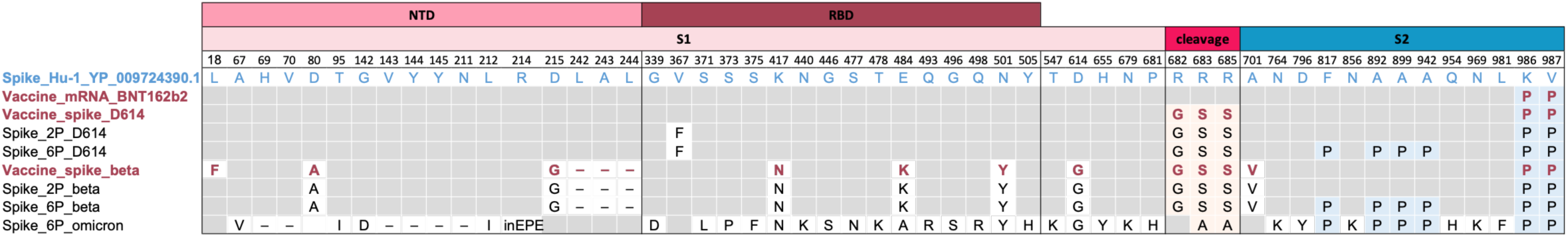
Spike sequence mutations across SARS-CoV-2 variants and stabilized constructs.

Ruling out increased overall immunogenicity of the MVB.1.351 vaccine, multiparametric flow cytometric analysis of the entire cohort at D28 or M3 revealed comparable quantitative recruitment of SARS-CoV-2 Hu-1 and/or B.1.351 spike-specific CD19^+^ IgD^-^ MBCs between groups (**Fig. 1c, S1a-b** and **Table S2a**) without selective expansion of B.1.351 spike monoreactive CD19^+^ IgD^-^ MBCs by the MVB.1.351 vaccine (**Fig. 1d** and **Table S2a**). As previously described by us and others in the context of limited antigenic drift between variants(Sokal et al. 2023; Alsoussi et al. 2023; Kaku et al. 2023), cross-reactive MBCs recognizing both ancestral Hu-1 and B.1.351 spike proteins represented approximately 80% of all spike-positive cells at any given time point post-boost in all groups (**Fig. 1d**), with overall similar frequencies of RBD-targeting cells among cross-reactive MBCs observed in both adjuvanted protein vaccine groups (**Fig. 1e** and **Table S2b**). The proportion of RBD-binding MBCs was transiently lower at D28 in the BNT162b2 vaccine group (**Fig. 1e**), but this effect was fully lost by the M3 time point. A similar picture could be seen at the serological level, with overall similar IgG titers against Hu-1 or B.1.351 RBDs in all groups, apart from a transient increase in Hu-1 RBD serum titers in the BNT162b2 vaccine group (**Table S1c**). While this highlights potential platform-related skewing of the MBC/antibody responses toward the RBD, it is unlikely to account for the superior neutralizing potency observed with MVB.1.351 as compared to both Hu-1 containing vaccines. Phenotypic analysis of CD19^+^IgD^-^ B cells further indicated comparable frequencies of CD71^+^ activated spike- and RBD-specific B cell across all three vaccine groups (**Fig. S1c-g** and **Table S2c-d**), although we cannot exclude potential differences at earlier time points as most RBD and spike-specific cells had already transitioned to a resting MBC phenotype by D28.

Focusing our analysis on the cross-reactive RBD-specific MBC response, we could distinguish more subtle qualitative differences specific to the MVB.1.351 group. In all individuals, two clear subsets, separated based on spike and RBD tetramer staining intensities, could be observed among RBD-specific Hu-1/B.1.351-cross-reactive CD19⁺IgD⁻ B cells (referred to here after as Beta^high^ and Beta^low^, **Fig. 1f** and **S1a-b**). This differential staining was especially marked for the B.1.351 spike (**Fig. 1f**) but also extended to all other spike variants included in our panel (**Fig. S1h**). Interestingly, these cells were not equally represented among vaccine groups. RBD-specific MBCs displaying an increased binding to the Beta B.1.351 spike trimer (Beta^high^, as opposed to the remaining Beta^low^ RBD-specific MBCs) appeared to be twice as frequent in MVB.1.351 boosted individuals at D28 and M3 compared to MVD614 and BNT162b2 vaccinees (**Fig. 1f-g** and **Table S2b**), reaching up to 40% of the RBD-specific compartment. This preferential expansion was not observed in individuals boosted with the MVD614 vaccine, indicating that this effect was not dependent of the vaccine platform or the adjuvant, but likely driven by the B.1.351 spike protein. Moreover, the frequency of Beta^high^ MBCs in individual donors correlated with the sera neutralizing activity against D614G, B.1.351 and BA.1 SARS-CoV-2 (**Fig. 1h**), suggesting a link between this Beta^high^ MBC expansion and the observed enhanced vaccine-induced protection in boosted individuals.

### Beta^high^ memory B cells are enriched in Hu-1/B.1.351 SARS-CoV-2 potent neutralizers

To assess the functionality of this RBD-specific Beta^high^ MBC population, we single-sorted Beta^high^ and Beta^low^ MBCs from 4 patients (3 from the MVB.1.351 group and 1 from the MVD614 group, **Fig. 2a**, and **Table S3a**), at early timepoints (D15 and D28 post-boost), prior to any likely substantial germinal center affinity maturation towards vaccine-specific epitopes. These MBCs were single-cell cultured and differentiated into antibody-secreting cells, followed by V(D)J sequencing of immunoglobulin heavy and light chains and direct functional testing of the monoclonal antibodies (mAbs) present in their culture supernatants (**Fig. 2a** and **Table S3a**) (Sokal, Barba-Spaeth, et al. 2021). In total, we were able to characterize 145 mAbs, including 82 from Beta^high^ MBCs and 63 from Beta^low^ MBCs (**Table S3a-b**).

**Fig. 2.**
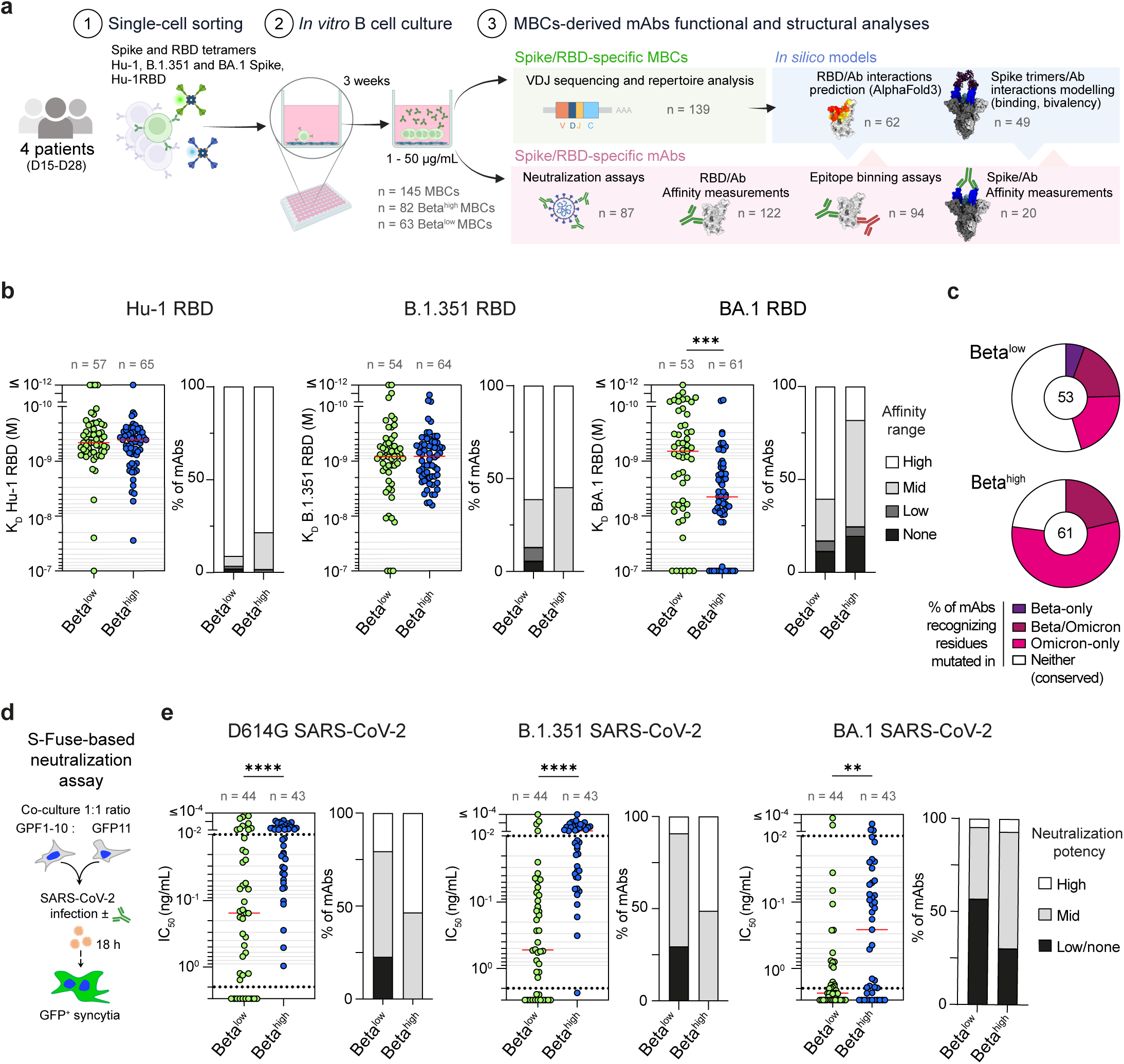
Beta^high^ memory B cells derived mAbs are potent neutralizers of D614G, B.1.351 and BA.1 SARS-CoV-2. (**a**) Overview of the experimental workflow for the functional and structural characterization of mAbs derived from Beta^high^ and Beta^low^ MBCs (single-cell culture supernatants). **(b)** Equilibrium dissociation constants (K_D_, in M) measured by BLI for Beta^low^ and Beta^high^ MBCs-derived mAbs binding to the Hu-1 (left panel), B.1.351 (middle panel) and BA.1 RBDs (right panel), with the corresponding frequencies of high-binders (K_D_ < 10^-9^ M), mid-binders (10^-9^ ≤ K_D_ < 10^-8^ M), low-binders (10^-8^ ≤ K_D_ < 10^-7^ M), and non-binders (K_D_ ≥ 10^-7^ M) displayed on the right side of each panel. (**c**) Pie chart representing the percentage of Beta^low^ and Beta^high^ mAbs inferred as recognizing RBD residues grouped as conserved between the ancestral strain B.1.351 and BA.1 (conserved, white), mutated only in B.1.351 (Beta only, mauve), mutated both in B.1.351 and BA.1 (Beta/Omicron shared, purple) or only mutated in BA.1 (Omicron Specific, pink). Number at the center of the pie-charts indicates the number of tested mAbs. MAbs were inferred as recognizing a given epitope group based on individual drop in K_D_ (ratio > 3) measured between the Hu-1 and the two tested variant RBDs. (**d**) Schematic view of the *in vitro* neutralization assay against authentic SARS-CoV-2 strains, using the S-Fuse neutralization assay. (**e**) *In vitro* neutralization half-maximal inhibitory concentration (IC_50_) and potency (high: IC_50_ < 0.01 ng/mL; mid: ≥ 0.01 ng/mL to < 2 ng/mL; low/none ≥ 2 ng/mL) of Beta^high^ and Beta^low^ MBC-derived mAbs against authentic D614G, B.1.351 and BA.1 SARS-CoV-2 strains. (b and e) Mann-Whitney test. \*\*\*\**p* < 0.0001, \*\*\**p* < 0.001, \*\**p* < 0.01, \**p* < 0.05. See also **Fig. S2** and **Table S3**.

Surprisingly, Beta^high^ and Beta^low^ MBC-derived mAbs displayed similar, and overall high, affinities towards Hu-1 RBD, in the nanomolar to sub-nanomolar range (**Fig. 2b, left**). Analysis of the underlying V(D)J sequences confirmed the recruitment of pre-existing germinal center-derived affinity-matured MBCs, with both Beta^high^ and Beta^low^ MBCs harboring similarly high numbers of immunoglobulin heavy chain variable region (IgV_H_) somatic hypermutations at these early time points (**Fig. S2a and Table S3c**). Repertoire analysis of both MBC subsets at the individual donor level revealed an increased frequency of expanded clones (**Fig. S2b**) and reduced Simpson’s diversity index (**Fig. S2c**) in Beta^high^ MBCs in all three donors with more than 10 sequences for each population, suggesting antigen-driven recruitment and expansion of specific clones in this population.

Ruling out specific recruitment of MBC bearing high affinity to specific epitopes in the B.1.351 spike included in the MVB.1.351 vaccine, Beta^high^ and Beta^low^ MBC-derived mAbs harbored similar affinities against both the Hu-1 and the B.1.351 RBD (**Fig. 2b, left and center**). A similar two-fold overall decrease in affinity to the B.1.351 RBD, compared to the Hu-1 RBD, could even be seen for both mAb pools (median: 4.1 x 10^-10^ and 8.15 x 10^-10^ respectively for Beta^high^ and 4.6 x 10^-10^ and 8.2 x 10^-10^ respectively for Beta^low^), with a clear loss in affinity (K_D_ ratio ≥ 3, see **Table S3d**) for 13/54 and 13/64 of mAbs in each group respectively. Affinity measurements towards the BA.1 RBD, however, revealed a specific impairment of Beta^high^ MBC-derived mAbs compared to Beta^low^ MBC-derived mAbs (**Fig. 2b, right**). Analyzing these results in the light of common and unique mutated residues in the B.1.351 and BA.1 RBD variants showed that both repertoires contained a similar frequency of mAbs sensitives to mutation found in both the B.1.351 and BA.1 RBD (namely K417N, E484K/A and N501Y); while the Beta^high^ repertoire appeared enriched in MBCs targeting residues mutated in Omicron BA.1, but not in former SARS-CoV-2 variants (**Fig. 2c**).

Notwithstanding comparable binding affinities, mAbs derived from Beta^high^ MBCs demonstrated markedly superior *in vitro* neutralization potency against both authentic D614G and B.1.351 SARS-CoV-2 (**Fig. 2d, e and Table S3g**), compared with those derived from Beta^low^ MBCs. More than half of the Beta^high^-derived mAbs exhibited IC_50_ values below 0.01 ng/mL, compared with only 20% and 9% of Beta^low^-derived mAbs against D614G and B.1.351 SARS-CoV-2, respectively (**Fig. 2e, left** and **center**). The same functional dichotomy between Beta^high^ and Beta^low^ mAbs was observed in individual vaccinated with the MVD614 vaccine, supporting the notion that the features of the Beta^high^ population are universal rather than vaccine specific (**Fig. S2d to f**). In line with the affinity data described above (**Fig. 2b**), the Beta^high^ mAb group displayed a strong reduction in *in vitro* neutralization potency against authentic BA.1 SARS-CoV-2. However, it remains important to note that Beta^high^ mAbs retained a significantly higher neutralization potency compared to their Beta^low^ counterparts (**Fig. 2e, right**). This is in line with the previously described impact of the MVB.1.351 vaccine at the serological level (Launay et al. 2022) and the observed correlation between Beta^high^ MBCs frequencies and circulating anti-BA.1 neutralizing Abs in boosted individuals (**Fig. 1h, right**).

Collectively, these findings are consistent with the targeted recruitment, expansion and maintenance of a subset of pre-existing cross-reactive RBD-specific MBCs by the MVB.1.351 vaccine. Their mAbs display potent SARS-CoV-2 neutralizing potency, correlate with the serological neutralizing response and can be predicted to target specific epitopes on the RBD based on their susceptibility to point mutations appearing in later variants. Our results also point towards the recruitment of Beta^high^ MBCs occurring independently of the direct recognition of B.1.351-specific neo-epitopes.

### Beta spike protein vaccine preferentially recruits class 1, 2 and 3 antibodies

Although the relatively limited number of sequences obtained constrained the type of in-depth repertoire analysis that could be performed, our V(D)J sequencing data did reveal differential IgV_H_ gene usage between Beta^high^ and Beta^low^ MBCs (**Fig. S2g** and **Table S3c**). On the one hand, V_H_3-66, frequently associated with public clonotypes of class 1 antibodies targeting the receptor binding motif (RBM) of the RBD was not detected in Beta^low^ MBCs and was instead enriched in Beta^high^ MBCs (**Fig. S2g)**(Barnes et al. 2020). One of these previously described public clonotype could further be detected in Beta^high^ MBCs of two out of the four donors analyzed (**Fig. S2h**)(Sokal, Chappert, et al. 2021; Nielsen et al. 2020). Similarly, V_H_1-58, V_H_2-5 and V_H_1-18, frequently used by class 1, 2 and 3 mAbs(Yuan et al. 2021), were also enriched in Beta^high^ MBCs. On the other hand, V_H_5-51, described to be associated with class 3, 4 and 5 mAbs(Starr et al. 2021), and V_H_3-23 were enriched in Beta^low^ compared to Beta^high^ MBCs (**Fig. S2g**).

We thus next applied two complementary approaches to map the epitopes of more than 90 mAbs present in our dataset (**Fig. 2a and Table S3a**): *in vitro* epitope-binning (EB) by biolayer interferometry on available culture supernatants (n = 94) (**Fig. S3a-c and Table S3f**) and parallel *in silico* modeling of mAb-RBD interactions for all paired Ig heavy and light chains (n = 133) using AlphaFold3(Abramson et al. 2024), an approach that we recently validated in the context of type-I IFN-specific mAbs(Fournier et al. 2026) (**Fig. S4, Table S4a-b**). EB experiments against a set of four reference mAbs covering the RBM (class 1 CB6 (PDB 7C01), class 2 LY-CoV555 (PDB 7KMG), class 3 REGN10987 (PDB 6XDG), and class 1/4 ADG2 (ADG20, PDB 7U2D) (**Fig. S3b**) revealed six distinct competition groups among the 94 tested culture supernatants (**Fig. 3a**). These initial experiments allowed clear separation of three class 1-competing groups (groups A to C) subdivided along the way of additional competition to representative class 2 and 1/4 mAbs. Two additional class 3-competing groups could be further identified (groups D and E) with one subgroup competing with class 1/4 ADG2 mAbs (group E) and another subgroup competing instead with class 2 mAbs (group D). The last group targeted epitopes away from the classical class 1, 2 and 3 epitopes and encompassed two additional reference mAbs targeting class 4 and 5 epitopes (group F).

**Fig. 3.**
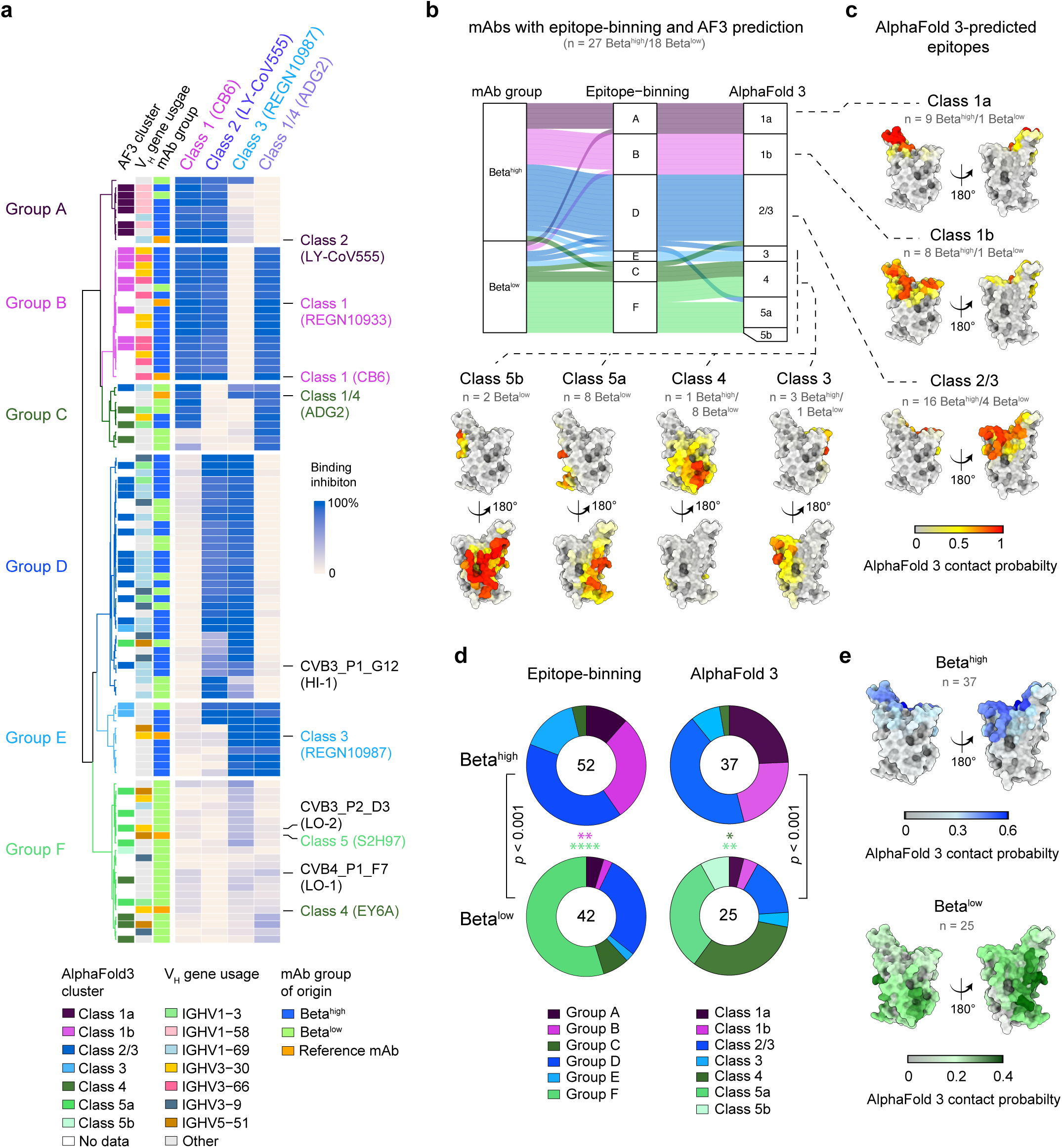
Beta^high^ and Beta^low^ MBCs recognize distinct RBD epitopes. (**a**) Epitope binning performed on 94 anti-RBD mAbs from MBC culture supernatants and 10 re-expressed anti-RBD mAbs (loaded first) against four reference mAbs (loaded second) using a competitive binding (BLI) sandwich assay with Hu-1 RBD. Data are expressed as percentage of binding inhibition: white denotes no inhibition, whereas dark-blue denotes complete inhibition. MAbs are hierarchically clustered based on their binding profiles (dendrogram). Row annotations indicate for each mAb the sorted group of origin (Beta^high^, Beta^low^, reference mAb), the IgV_H_ gene usage, for the most represented IgV_H_ (n ≥ 5) and the epitope group defined by AlphaFold3 modeling when available. (**b**) Sankey diagram illustrating the relationships between RBD antibody classes determined by epitope-binning (center), and epitope groups derived from AlphaFold3 mAb-Hu-1 RBD interaction predictions (left) for all tested and confidently predicted Beta^high^ (top) and Beta^low^-MBC derived mAbs (bottom). (**c**) Average AlphaFold3-predicted contact probabilities projected on the Hu-1 RBD structure for mAbs of indicated AlphaFold3 epitope clusters. (**d**) Pie-charts representing the distribution of RBD-specific Beta^high^ (top) and Beta^low^ (bottom)-derived mAbs among epitope-binning groups (left) or AlphaFold3-predicted classes (right). (**e**) Average AlphaFold3-predicted contact probabilities projected on the Hu-1 RBD structure for Beta^high^ (top, blue) and Beta^low^ (bottom, green)-derived mAbs. (d and e) Global Chi-square test before individual Fisher’s test with Bonferroni correction for each class. Global *p*-value is indicated in black and significant comparison for each category are color-coded. See also **Fig. S3, S4**, **Tables S3 and S4**.

Parallel *in silico* modeling of mAb/Hu-1 RBD interactions for 133 IgH/IgL pairs, either as variable fragments (Fvs) or antigen-binding fragments (Fabs) regions, provided us 62 confident predictions (see **Methods, Fig. S4a-B** and **Table S4a)**, including 45 of the 94 EB tested mAbs. While we cannot exclude limited IgV_H_ and epitope-based bias in the overall quality of AlphaFold3 predictions (**Fig. S4c**-**d**), unsupervised clustering of predicted binding patterns for all confident *in silico* models generated 7 clusters of binding patterns mostly overlapping with the groups identified by epitope-binning (**Fig. 3b** to **d and S4e,** and **Table S4b**). Of interest, these predictions allowed us to better delineate the epitopes recognized by EB group F members, which could now be classified into three distinct classes: 4, 5a, and 5b (**Fig. 3b** to **d**). Class 4 mAbs targeted the same epitope region as those from EB group C, albeit with different binding angles (**Fig. S3d to g**). While, as previously described(Starr et al. 2021), classes 5a and 5b targeted two slightly overlapping additional epitopes on the opposite side of bottom of the RBD (**Fig. 3b** to **d**). These predictions were further validated by expressing one mAb member for each classes 2/3, 4 and 5a (class 2/3: HI-1, class 4: LO-1 and class 5a: LO-2) and performing reciprocal EB assays with additional reference mAbs targeting class 4 and 5 epitopes (class 4 EY6A (PDB 6ZER), class 5 S2H97 (PDB 7M7W), **Fig. S3c** and **Table S3f**).

Overall, Beta^high^ MBC-derived mAbs appeared strongly enriched in class 1a, 1b and 3 (n = 29/52 (55.7%) of mAbs in EB assays and n = 20/37 (54%) of mAbs confidently predicted by AlphaFold3 (AF3)) (**Fig. 3d**) and were, as a whole, mostly predicted to target epitopes overlapping with the RBM at the top of the RBD (**Fig. 3e**). In contrast, Beta^low^ MBC-derived mAbs showed preferential binding towards class 4 and 5 epitopes (n = 26/42 (61.9%) in EB assays and n = 18/25 (72%) in AF3 predictions) (**Fig. 3d**), mostly positioned at the base of the RBD (**Fig. 3e**), with both classes of mAbs being largely absent from Beta^high^ MBC-derived mAbs (n = 2/52 (3.8%) in EB assays and n = 1/37 (2.7%) in AF3 predictions). While class 4 and 5 epitopes are highly conserved(Starr et al. 2021; Jette et al. 2021; Burnett et al. 2021; Jensen et al. 2023; Cui et al. 2024), they display limited overlap with the RBM and are predicted to be mostly buried in most conformational states of the pre-fusion trimer of the SARS-CoV-2 spike. This correlated with lower potency to neutralize D614G, B.1.351 and BA.1 SARS-CoV-2 strains compared to all other classes (**Fig. S3h**), providing a mechanistical explanation to the observed difference in neutralization potency between Beta^high^ and Beta^low^ MBC-derived mAbs (**Fig. 2e**).

### Beta^high^ and Beta^low^ antibodies exhibit different binding properties to the canonical spike trimer

The structural insights gained from competitive binding experiments and *in silico* modeling raised two potential, and not mutually exclusive, hypotheses to explain the differential binding to tetramers of spike trimers for Beta^high^ and Beta^low^ MBCs: 1/ differential epitope accessibility in the context of the spike trimers or 2/ bivalent binding to two RBDs within the same spike trimer, or intra-spike crosslinking, resulting in increased overall avidity as suggested for some class 1 mAbs(Barnes et al. 2020). To distinguish between these two hypotheses, we first set up an *in-silico* workflow to assess the binding properties of our mAbs across multiple experimentally resolved SARS-CoV-2 spike trimer structures. Confident Fab/Hu-1 RBD AlphaFold3-predicted models (n = 49) were aligned to canonical spike trimers with zero (PDB 7BNM), one (PDB 7BNN) or two (PDB 7BNO) RBDs in the “up” conformation(Benton et al. 2021), as well as an expanded open trimer conformation stabilized upon antibody binding (S304 mAb, PDB 7JW0(Piccoli et al. 2020)), enabling the systematic assessment of steric clashes for individual Fab-spike complexes, as well as inter-Fab distances as representative for epitope accessibility and bivalent engagement respectively (**Fig. 4a**).

**Fig. 4.**
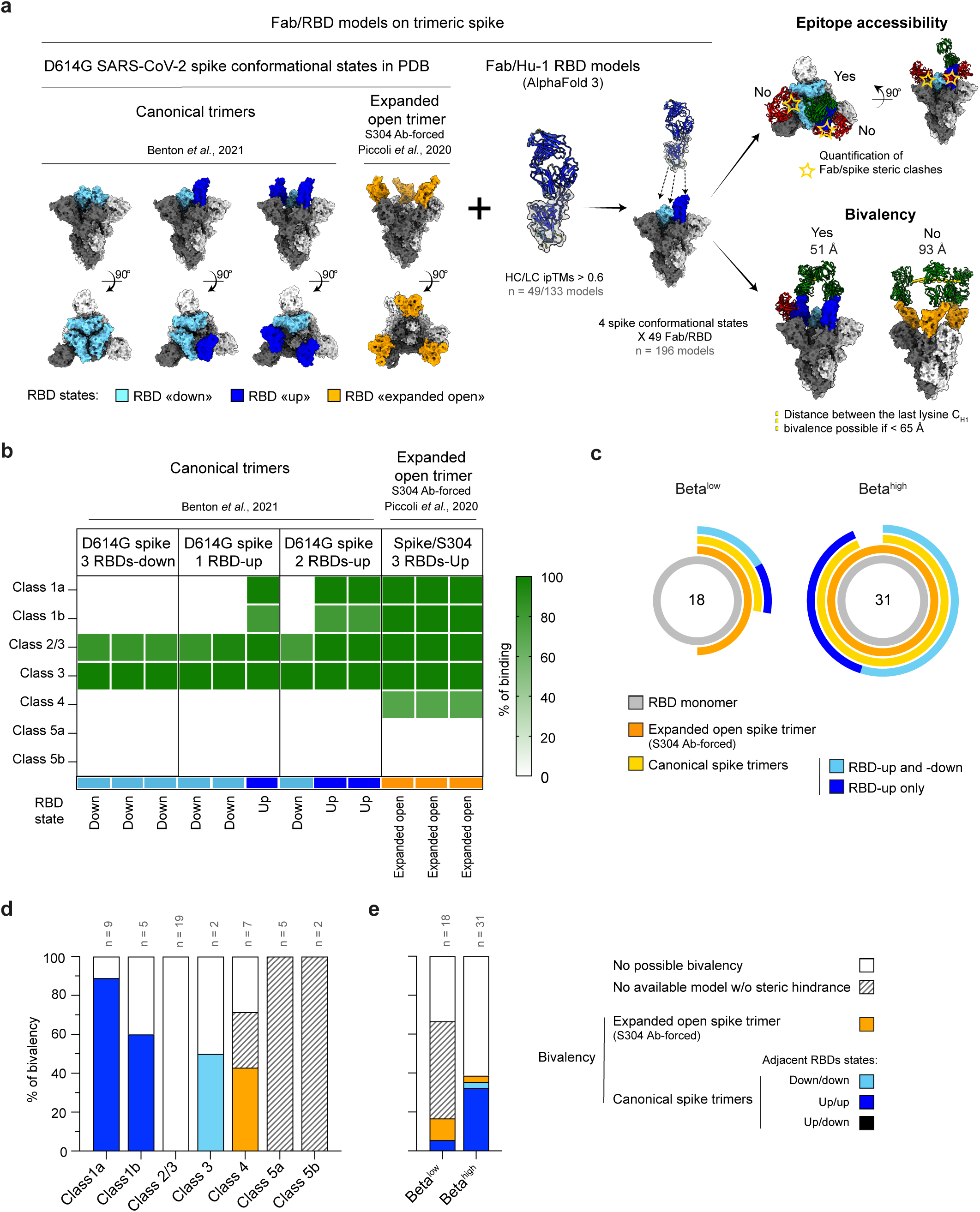
Beta^high^ recognize accessible spike epitopes, while Beta^low^ are enriched in cryptic spike epitopes. (**a**) Overview of the *in-silico* pipeline used to assess binding properties of anti-RBD mAbs in the context of the full D614G SARS-CoV-2 prefusion spike trimer. AlphaFold3-predicted Fab/RBD complexes with confident predictions scores (**see Methods**) were first aligned onto four cryo-EM-resolved conformational states of D614G SARS-CoV-2 spike protein: the canonical trimeric conformation displaying zero, one, or two receptor-binding domains (RBDs) in the “up” state (PDB 7BNM, 7BNN, 7BNO), and an expanded open trimer stabilized upon antibody binding (S304, PDB 7JW0). Epitope accessibility (no detectable steric clash of bound Fab with residues outside of the bound RBD) and Ab bivalency (distance between the carbons of the last lysine of two bound Fab C_H1_ domains < 65Å) were subsequently quantified. In canonical spike trimers, RBDs-up are shown in dark blue, whereas RBDs-down are depicted in light blue. In the expanded open trimer, RBDs are shown in yellow. To illustrate the diversity of binding configurations, 3 Fabs of a representative class 1a mAb (CVB1_P6_G5) are shown aligned onto each protomer of selected spike conformational states. Fabs are colored green when no steric clash could be detected, and red otherwise. (**b**) Heatmap depicting the proportion of assessed mAbs, stratified by epitope class, predicted to bind each individual RBD of an indicated conformational state of the prefusion spike trimer without steric clashes without any of clash with the rest of the trimer. (**c**) Pie charts showing the frequencies of assessed mAbs, stratified by sorted MBC subset (Beta^high^ or Beta^low^) predicted to bind to indicated conformational state of the prefusion spike trimer without any steric clash with the rest of the trimer. The central value refers to the total number of mAbs analyzed. (**d** and **e**) Percentage of mAbs, stratified by AlphaFold3-predicted class (d) or sorted MBC subset (Beta^high^ or Beta^low^), predicted to be able to bind bivalently. White bars represent no possible bivalent binding, colored bar represent predicted bivalent binding to two RBDs, with colors indicating the RBD/spike conformation permitting bivalency. Hatched areas indicate mAbs from class 4, 5a and 5b for which bivalency could not be evaluated due to the lack of a prefusion spike conformation allowing binding without steric hindrance. See also **Fig. S5** and **Table S4**.

Our analysis confirmed that most class 2/3 and 3 mAbs identified in our dataset, regardless of their MBC of origin, are predicted to reach their epitopes without major clash in the context of all canonical D614G spike trimer structures (**Fig. 4b, Table S4c**), in line with previously published X-ray structures(Qi et al. 2022; Barnes et al. 2020). Most class 1a and 1b are also predicted to bind the canonical D614G spike trimer but only to bind RBDs in their “up” state in the complex (**Fig. 4b and S5a**). In contrast, all epitopes targeted by class 4 or 5 mAbs in our dataset appeared buried inside the canonical spike trimer (**Fig. 4b and S5a**). Class 4 mAbs were only predicted to reach their epitope in the context of S304-stabilized expanded opened trimer conformation (PDB 7JW0(Piccoli et al. 2020), **Fig. 4b and S5a**). To our knowledge, no comparable structure has been published for a class 5a or 5b mAb, likely due to the significant displacement of the N-terminal domain (NTD) required to access their antigenic site(Y. Luo et al. 2026). Consistently, we were unable to model sterically compatible binding of these two subclasses to a spike trimer (**Fig. 4b and S5a**). Nonetheless, analyzing these data along the lines of MBC groups confirmed that Beta^high^ and Beta^low^ MBC-derivedmAbs strongly differed in their ability to bind the canonical spike trimer (**Fig. 4c**), with over 90% (29/31) Beta^high^ MBC-derived mAbs predicted to bind the canonical D614G spike trimer without clash when only a third (5/18) of Beta^low^ MBC-derived mAbs were able to bind a spike trimer without forcing any important conformational changes.

Based on these alignments, inter-Fabs distance could further be calculated for all mAbs, in all contexts in which two Fabs could bind to the same trimer without significant steric hindrance (**Fig. S5b, Table S4c**). When doing so, most of the class 1 mAbs were predicted to bind bivalently to two RBDs in the “up” state on the spike trimer (**Fig. 4d** and **S5b**). Bivalency was also predicted for half of class 3 mAbs, when bound to two RBDs in the down conformation, as well as up to 40% of class 4 mAbs, when bound to RBDs in the expanded open spike trimer. In contrast, bivalency appeared fully precluded for class 2/3 mAbs due to individual Fabs facing away from one another when bound to the spike trimer (**Fig. 4d** and **S5b**). Overall, bivalency could only be predicted for up to 40% of Beta^high^ MBCs and less than 20% of Beta^low^ MBCs (**Fig. 4e**). A role for bivalency in the increased neutralization potential of some of the class 1 and class 3 Beta^high^ MBC-derived mAbs can be considered, as previously demonstrated(Callaway et al. 2023). Bivalency, however, is unlikely to explain the enhanced recruitment of most of Beta^high^ MBCs, in contrast with the clear difference in availability described above (**Fig. 4c**).

### MVB.1.351 vaccine restrains response against the inner RBD epitopes via its conformational properties

To experimentally validate the *in silico*-predicted differences in epitope accessibility, we next quantified the binding affinities of mAbs derived from Beta^high^ and Beta^low^ MBCs to Hu-1 and B.1.351 spike trimers using surface plasmon resonance (SPR). Notably, we employed here 6-proline-stabilized spike trimers (S-6P), as used in our initial tetramer staining strategy, to first link SPR affinity measurements with FACS staining used to stratify mAbs according to their MBC origin (Beta^high^, Beta^low^). To separate differences in epitope accessibility in the context of spike trimers from additional variation in recognition due to variant-specific epitope mutations, we also restricted this analysis to mAbs exhibiting comparable affinities for Hu-1 and B.1.351 RBDs (K_D_ ratio between 0.5 and 1.5, n = 21). We further randomly sampled from this subset (**Fig. S6a** and **Table S3e**), blinded to mAb classes. Consistent with *in silico* structural predictions, Beta^low^ MBC-derived mAbs displayed strikingly reduced binding affinities to both Hu-1 and B.1.351 S-6P as compared to their Beta^high^ MBC-derived counterparts (**Fig. 5a** and **S6b**). This effect was primarily driven by reduced association rates (k_on_) (**Fig. 5b** and **S6c**), in line with the predicted requirement for a destabilized spike conformation to enable binding to cryptic epitopes frequently targeted by Beta^low^ MBC-derived mAbs. Interestingly, recapitulating the observed small difference in tetramer staining (**Fig. S1h**), association-rates were even more reduced for Beta^low^ MBC-derived mAbs in the context of the B.1.351 S-6P compared to the Hu-1 S-6P (**Fig. 5c**).

**Fig. 5.**
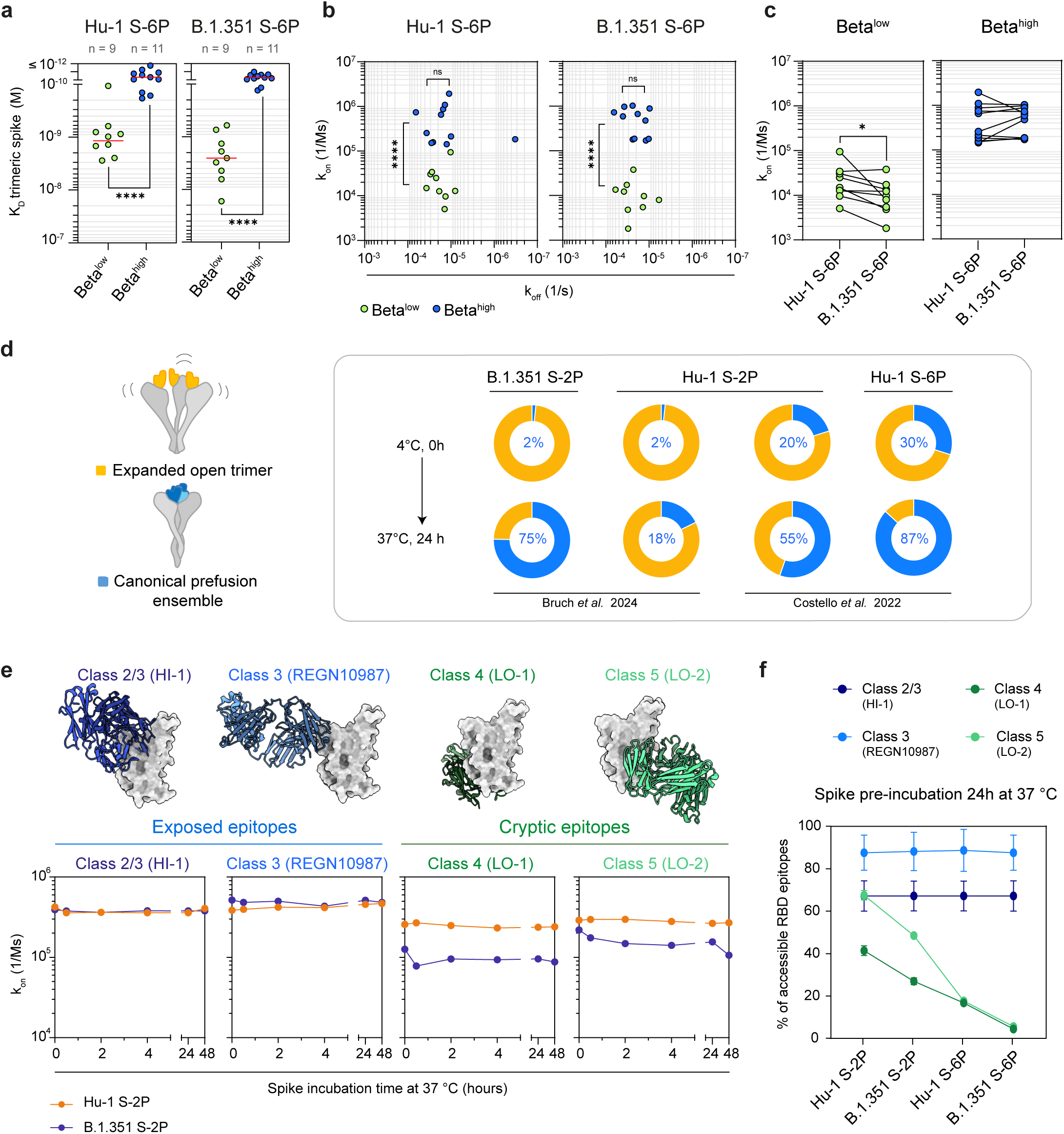
B.1.351 spike conformational determinants restrain access to cryptic B cell epitopes, favoring the recruitment of Beta^high^ MBCs. (**a-c**) Equilibrium dissociation constants (K_D_ = k_on_/k_off_, in M) (A), scatter plot of association (k_on_, 1/Ms) vs dissociation (k_off_, 1/s) rate constants (b) and paired-association rate constants (k_on_, 1/Ms) (c) measured by Surface Plasmon Resonance (SPR) for Beta^low^ (n = 9, green) and Beta^high^ (n = 11, blue) MBCs-derived mAbs binding to the 6 prolines-stabilized (S-6P) Hu-1 (left) or B.1.351 spike trimers (right). Tested mAbs were randomly selected among mAbs displaying similar binding affinities (**see** Fig. 2b) to the Hu-1 and B.1.351 RBDs in both groups. (**d**) Published relative proportions of expanded open (yellow) and canonical spike trimer dynamics states (blue) in 2 prolines-stabilized (S-2P) or 6 prolines-stabilized (S-6P) Hu-1 or B.1.351 S-2P following 24 h incubation at 37 °C(Bruch et al. 2024; Costello et al. 2022). (**e**) Time-dependent changes of apparent k_on_ (1/Ms) measured by BLI, for four re-expressed representative mAbs, targeting either exposed epitopes (HI-1 - predicted class 2/3 (CVB3_P1_G12) from our dataset - and class 3 REGN10987(Hansen et al. 2020)) or cryptic epitopes LO-1 - predicted class 4 (CVB4_P1_F7) - and LO-2 - predicted class 5 (CVB3_P2_D3) - from our dataset) tested against 4 °C-stored Hu-1 and B.1.351 S-2P that had subsequently been incubated at 37 °C for the indicated number of hours. (**f**) Estimated accessibility for indicated RBD epitopes-specific mAbs (HI-1, REGN10987, LO-1 and LO-2) after 24 h incubation of indicated prefusion spike trimer at 37 °C. For each mAb/trimer pair, accessibility (percent of maximum accessibility) was calculated by estimating the ligand concentration correction required for measured apparent k_on_ against indicated spike trimer to equate true k_on_ measured against an isolated RBD monomer of the same variant (**see Methods**). Experiments were performed in triplicate. (a and c) Mann-Whitney test. \*\*\*\**p* < 0.0001, \*\*\**p* < 0.001, \*\**p* < 0.01, \**p* < 0.05. See also **Fig. S6** and **Table S3**.

The structure of both Hu-1 and B.1.351 spike trimers contained in the MVD614 and the MVB.1.351 vaccines has been resolved(Bruch et al. 2024), pointing to two main differences: 1/ the spike trimer in the MVB.1.351 vaccine preferentially adopts the canonical conformational state with two RBDs-up, favoring bivalent engagement of class 1 anti-RBD mAbs, and 2/ the reversible equilibrium between canonical and expanded open trimer states shifts toward the canonical trimer state at 37 °C for the MVB.1.351, as compared to the MVD614. This is consistent with a slower transition from the expanded open trimer, favored at 4 °C, to the canonical trimer state (**Fig. 5d)**(Bruch et al. 2024). This equilibrium is further shifted toward the canonical state in the 6-proline-stabilized spike trimer (**Fig. 5d)**(Costello et al. 2022). Because bivalent binding of class 1 mAbs present among Beta^high^ MBC-derived mAbs could not fully account for the enhanced recruitment of Beta^high^ MBCs upon boosting with the MVB.1.351 vaccine, we next investigated whether differences in spike trimer conformational change from cold storage to body temperature, translating in differential exposure of RBD epitopes, may contribute to this preferential recruitment.

Our SPR-measurements already suggested reduced accessibility of class 4 and 5 cryptic epitopes in the context of the B.1.351 spike (**Fig. 5c**); however, this experiment was performed at 25 °C using a 6-proline stabilized spike and therefore only partially recapitulated physiological conditions that occur early after vaccination. Upon boosting with the MVB.1.351 and MVD614 vaccines, donor’s immune system likely encountered a 2-proline stabilized spike (S-2P) at body temperature and most likely still in the process of transitioning from 4 °C storage. To better delineate epitope accessibility in such context, we assessed the binding capacity of representative reference mAbs (class 3 REGN10987) or re-expressed Beta^high^ (class 2/3 CVB3_P1_G12, hereafter referred to as HI-1) and Beta^low^ MBC-derived mAbs (class 4 CVB4_P1_F7 and class 5a CVB3_P2_D3, hereafter referred to as LO-1 and LO-2 respectively) to multiple spike trimers following transfer from 4°C to 37°C for various incubation times using biolayer interferometry (**Fig. 5e** and **f**). In the context of S-2P, representative mAbs derived from Beta^high^ MBCs displayed comparable and stable association rates over time (k_on_, 1/Ms) for both Hu-1 and B.1.351 spikes. In contrast, Beta^low^ MBC-derived representative mAbs showed a rapid decline in measured association rates within the first 30 minutes of incubation at 37 °C for the B.1.351 spike (**Fig. 5e**), but not the Hu-1 spike, in line with the more rapid conformation shift observed by Bruch and colleagues for the B.1.351 spike trimer(Bruch et al. 2024). These reduced k_on_ values then persisted for up to 48 h. Repeating these experiments to compare S-2P and S-6P, and converting measured association rates into estimated percentages of accessible RBD epitopes (see **Methods**) for each mAb after 24 h at 37 °C, further revealed a synergistic effect between B.1.351-specific mutations and additional proline-stabilization in drastically restricting access to cryptic class 4 and 5 epitopes (**Fig. 5f**).

Overall, our results demonstrate that the conformational dynamics of the B.1.351 spike trimer restrains access to cryptic epitopes, favoring recall response against exposed RBD epitopes, explaining the stronger and broader elicited neutralizing responses. Our results also suggest that the recruitment of Beta^high^ MBCs could be even more pronounced with an S-6P, as previous results have already suggested(Costello et al. 2022; Bruch et al. 2024; Stuible et al. 2023).

## DISCUSSION

The efficacy and speed of design of current SARS-CoV-2 vaccines have drawn immensely on prior knowledge regarding key mutations required to stabilize class I fusion proteins in their pre-fusion form, knowledge mostly derived from earlier works on HIV, RSV, and other coronaviruses(Sanders and Moore 2021). Our results suggest that additional mutations, naturally selected in viral variants, exert a synergistic effect with these stabilizing mutations to refocus the humoral response towards neutralizing epitopes at the top of the RBD and may be incorporated to the “reverse vaccinology” toolbox. More importantly, we provide an *in vivo* demonstration that subtle differences in the conformational dynamics of the spike can induce a rapid and durable remodeling of an existing human MBC repertoire towards such epitopes.

Stabilization of the spike trimer through proline substitutions influences its structure and immunogenicity(Corbett et al. 2020). Several studies using different vaccine platforms in mice and non-human primates have consistently shown that the S-2P elicits stronger immune responses than its wild-type counterpart(Mercado et al. 2020; Bos et al. 2020; Kalnin et al. 2021). Accordingly, most spike vaccines approved as of mid-2026 in the European Union and North America, including those developed by Pfizer-BioNTech, Moderna and Novavax, incorporate these two-proline substitutions and most also include additional modifications at the cleavage site (RRAR to GSAS). These mutations stabilize the spike trimer in its prefusion state, reducing its conformational dynamics. Additional SARS-CoV-2 variant-specific mutations affect the reversible equilibrium observed between various canonical and the expanded open-trimer state of the prefusion spike trimer(Costello et al. 2022; Stuible et al. 2023; Edwards et al. 2021). These conformational variants likely impact how conserved epitopes buried within the prefusion conformation, notably underneath the RBD in the “down” position(Benton et al. 2020; Wrapp et al. 2020), are exposed to the immune system. Our results confirm such predictions by showing that even a small shift in epitope accessibility (an estimated 15% drop in accessibility for cryptic class 4 and 5 epitopes between the spike trimers included in the MVD614 and the MVB.1.351 vaccines, respectively) appeared sufficient to promote significant refocusing of the B cell responses toward immunodominant epitopes at the top of the RBD in the context of the COVIBOOST trial (from 20 to 40% of the overall repertoire). Outside the RBD and the NTD, spike proteins from the MVD614 and the MVB.1.351 vaccines differ only by D614G and A701V, the latter being absent from Omicron lineages (**Table 1**)(Uraki et al. 2026; Bruch et al. 2024). Additional studies are warranted to determine whether incorporating these particular mutations in the design of future variant-targeted vaccines could provide a similar benefit by favoring the canonical conformation with two RBDs in the “up” state(Bruch et al. 2024). Conversely, our results suggest that the impact of variant-associated mutations on conformational preference should be monitored when designing variant-targeted vaccines to remove mutations that favor the dynamics toward the open trimer.

Most initial assessments of the impact of prefusion stabilization on the immunogenicity of the spike protein have been done in the context of primary responses(Mercado et al. 2020; Bos et al. 2020; Kalnin et al. 2021). A key question remained whether conformational changes of the spike protein might overcome immune imprinting. Our results demonstrate the selective mobilization of a subset of the MBC compartment displaying enhanced neutralization potential against B.1.351 SARS-CoV-2 as well as against both the D614G and the more distant Omicron BA.1 strains.

This resulted in a rapid and sustained remodeling of the MBC pool over time, independently of the recruitment of naive B cells targeting mutated epitopes within the B.1.351 spike, a point already documented in both vaccination and infection contexts(Kaku et al. 2023; Sokal et al. 2023). The parallel increase in serum neutralizing titers after MVB.1.351 vaccination(Launay et al. 2022) likely reflects the differentiation of part of these mobilized MBCs into plasma cells, and these titers also appeared stable up to three months.

Beside inducing high and stable neutralizing potency towards the vaccine strain, a central challenge in vaccine design remains in achieving durable breadth, so that immune responses remain effective against evolving SARS-CoV-2 variants. While epitopes on the top of the RBD elicit highly potent neutralizing responses, they are also primary sites for viral escape mutations(Starr et al. 2021). Conversely, cryptic epitopes within the inner RBD are less frequently neutralizing but remain highly conserved across SARS-CoV-2 lineages, serving as critical targets for broadly neutralizing antibodies(Starr et al. 2021; Jette et al. 2021; Burnett et al. 2021; Jensen et al. 2023; Cui et al. 2024). Our results highlight a fundamental trade-off between neutralizing potency and epitope breadth imposed by the spike trimer dynamics. From a vaccine perspective, strategies could be tailored. Highly stabilized canonical spike trimer could be used to preferentially recruit potent neutralizing MBCs and induce rapid protection against emerging SARS-CoV-2 variants in pre-immunized individuals, circumventing prior immune imprinting(Lu et al. 2022; Launay et al. 2022). Instead, in a context of prime immunization and pandemic preparedness, one could favor less stabilized spike-based strategies or subdomain-based immunogens predicted to induce a broader and more diverse antibody repertoire, including a larger share of antibodies targeting conserved epitopes on the pre-fusion spike trimer across sarbecoviruses. Recent studies in this direction suggest that non-stabilized spike immunization indeed redirects antibody responses toward conserved epitopes on the inner face of the RBD(Malewana et al. 2025). These findings should be interpreted with caution. Decades of research on the RSV class I fusion protein have underscored the critical importance of a minimal stabilization of the prefusion state to maintain a proper balance for the humoral response between the pre- and the post-fusion forms of the F trimer, and notably preserve sufficient targeting of neutralizing antigenic sites Ø and V that are otherwise not present in the post-fusion form(Ruckwardt et al. 2019). In this regard, vaccine strategies based on isolated RBD and NTD regions such as the recently approved mRNA-1283 may help further enhance the accessibility of RBD cryptic epitopes while limiting non-neutralizing responses against the spike S2 domain(Chalkias et al. 2025).

Overall, our findings reinforce the key impact of spike trimer antigen conformational dynamics in the remodeling of the B cell repertoire that occurs upon vaccine boost, with both short- and long-term effects on protection. Such impact should be carefully monitored in future clinical trials and incorporated in the rational design of future spike-based vaccines.

## MATERIALS AND METHODS

### Study design and human subjects

Subjects included in this study were part of the COVIBOOST trial(Launay et al. 2022). COVIBOOST was a randomized, single-blinded, multicenter trial across 11 centers in France. Participants were recruited from December 8, 2021, to January 14, 2022. The protocol was conducted in accordance with the Declaration of Helsinki and French law for biomedical research. It was approved by the “CPP Ile de France III” Ethics Committee and the French Health Products Safety Agency (ANSM). The study is registered with the ClinicalTrials.gov identifier NCT05124171 and with the EudraCT identifier 2021-004550-33. All donor-related information is listed in **Table S1**.

Median age was 45 years old (range 19-70 years old). These subjects had previously received two doses of BNT162b2 mRNA vaccine (ancestral strain) at least 5 months (median: 178 days, range: 151-206 days) before and were mostly COVID-19 naïve, except one subject in the MVD614 group who had a positive IgG nucleocapsid serology from baseline (**Table S1**)(Launay et al. 2022). Included subjects had a stable medical condition and no obvious immunosuppression condition that could alter significantly vaccine response. Subjects included were randomized to receive a booster dose of recombinant protein-based subunit vaccine or mRNA vaccine. The mRNA vaccine was a homologous booster using the BNT162b2 mRNA vaccines encoding the ancestral strain (Comirnaty, Pfizer BioNTech). The protein-based vaccines were either containing recombinant spike of the ancestral strain (MVD614, Sanofi-Pasteur/GSK), or of the Beta (B.1.351) strain (MVB.1.351, VidPrevtyn, Sanofi-Pasteur/GSK). Both were adjuvanted with AS03 adjuvant (squalene-based adjuvant). Participants were sampled at 15, 28 days and 3 months post-booster vaccination. While some patients had a nucleocapsid IgG seroconversion during the follow-up, no MBCs were functionally characterized after seroconversion. PBMCs were isolated from venous blood samples *via* standard density gradient centrifugation and cryopreserved at - 150 °C.

### Neutralization assays on sera

Neutralizing serum titers presented as part of Fig. 1b were directly extracted from the original publication of the COVIBOOST trial clinical results(Launay et al. 2022), for which methods have been reported. Sera neutralization potential of Wuhan (D614), Beta (B.1.351) and Omicron (BA.1) SARS-CoV-2 were assessed using a microneutralization test. The test uses clinical strains of SARS-CoV-2 (100 TCID_50_/well), TMPRSS2-expressing VeroE6 cells and relies on cytopathic effect (CPE) identification at 5 days post-infection. It is a VNT100 (100% of wells lysed in duplicate format). The test is automated in a NSB3 laboratory for all dilution and dispensing steps and for CPE reading. Dilutions tested were 20, 40, 80, 160, 320, 640 and 1280. The range was extended if a titer of 1280 was observed in the first instance.

### S-Fuse neutralization assay

U2OS-ACE2 GFP1-10 or GFP 11 cells, also termed S-Fuse cells, become GFP^+^ when they are productively infected by SARS-CoV-2(Buchrieser et al. 2020; Planas et al. 2021). Cells tested negative for mycoplasma. Cells were mixed (ratio 1:1) and plated at 8 × 10^3^ per well in a μClear 96-well plate (Greiner Bio-One). The indicated SARS-CoV-2 strains were incubated with serially diluted single MBC culture supernatants for 15 min at room temperature and added to S-Fuse cells. 18 h later, cells were fixed with 4% PFA (Electron Microscopy Sciences, # 15714-S), washed and stained with Hoechst (dilution of 1:10,000, Invitrogen, # H3570). Images were acquired using an Opera Phenix high-content confocal microscope (PerkinElmer). The GFP area and the number of nuclei were quantified using the Harmony software (PerkinElmer). The number of GFP syncytia and the number of nuclei were quantified using Harmony software (PerkinElmer). The percentage of neutralization was calculated using the number of syncytia as value with the following formula: 100 × (1 – [value with supernatant – value in “non-infected”]/[value in “no supernatant” – value in “non-infected”]). For each supernatant, the half-maximal inhibitory concentration (IC_50_, in ng/mL) was calculated with a reconstructed curve using the percentage of neutralization at each concentration. Neutralization potency of supernatants was classified as “high” (< 0.01 ng/mL), “mid” (≥ 0.01 ng/mL to < 2 ng/mL), and “low/none” (≥ 2 ng/mL).

### Virus strains

The reference strain D614G (hCoV-19/France/GES-1973/2020) was provided by the National Reference Centre for Respiratory Viruses hosted by Institut Pasteur (Paris, France) under the direction of Professor. S. Van der Werf. This viral strain was procured via European Virus Archive goes Global (Evag) platform, which has been supported by funding from the European Union’s Horizon 2020 research and innovation program under grant agreement # 653316. The Beta (B.1.351) variant (CNR 202100078) was first identified in an individual residing in Créteil, France(Planas et al. 2021). The Omicron variants BA.1 (hCoV-19/Belgium/rega-20174/2021) was provided and sequenced by the NRC UZ/KU Leuven in Leuven, Belgium(Planas et al. 2022).

Informed consent was provided by all patients or legal representatives for the utilization of biological materials. Individuals provided informed consent for the use of their biological materials. The variant strains were isolated from nasal swabs on Vero E6 or IGROV-1 cells and amplified by one or two passages on Vero cells.

Titration of viral stocks was performed on Vero E6 cells, with a limiting dilution technique allowing a calculation of the 50% tissue culture infectious dose, or on S-Fuse cells. Viruses were sequenced directly on nasal swabs and after one or two passages on Vero E6 or IGROV-1 cells. The sequences were deposited on GISAID immediately following their generation (D614G: EPI_ISL_414631; Beta: [CNR 202100078]; Omicron BA.1 ID: EPI_ISL_6794907.)

### Recombinant antigen production and purification

#### Construct design

Genes coding for SARS-CoV-2 spike ectodomains (Hu-1 and B.1.351) with Hisx8, Strep and Avi tags were synthesized by Genscript and cloned into the pcDNA3.1(+) vector. The ectodomains (residues 1-1208, Hu-1 numbering) were stabilized to preserve their trimeric prefusion conformation by introducing two (K986P, V987P, Hu-1 numbering) or six proline substitutions (F817P, A892P, A899P, A942P, K986P, V987P, Hu-1 numbering), a GSAS substitution at the furin cleavage site (residues 682–685) and a C-terminal Foldon trimerization motif(Hsieh et al. 2020). The S2P-stabilized constructs have the K986P and V987P substitutions, the GSAS linker instead of the furin site and the Foldon motif.

The SARS-CoV-2 Hu-1, B.1.351 and BA.1 RBDs were cloned in pcDNA3.1(+) encompassing residues 331-528 (Hu-1 numbering) from the spike ectodomains, and they were flanked by an N-terminal IgK signal peptide and a C-terminal Thrombin cleavage site followed by Hisx8-Strep-Avi tags in tandem.

#### Protein expression and purification

The plasmids coding for the recombinant proteins were transiently transfected in Expi293F™ cells (Thermo Fischer Scientific) using FectroPRO^®^ DNA transfection reagent (Polyplus), according to the manufacturer’s instructions. The cells were incubated at 37 °C (Hu-1 S, B.1.351 S, Hu-1 RBD, B.1.351 RBD) or 32 °C (BA.1 RBD) for 5 days and then the culture was centrifuged, and the supernatant was concentrated. The proteins were purified from the supernatant by affinity chromatography on a StrepTactin column (IBA). The spike proteins were further purified by size-exclusion chromatography (SEC) on a Superose6 increase 10/300 column (Cytiva) equilibrated in PBS, while the RBDs were loaded onto a Superdex200 increase 10/300 column (Cytiva).

#### Protein biotinylation

Hu-1 RBD, Hu-1 S-6P and B.1.351 S-6P Avi-tagged proteins were biotinylated using the Avidity BirA biotin-protein ligase kit according to the manufacturer’s instructions. Bovine serum albumin was biotinylated using EZ link NHS biotin (Thermo Fischer Scientific) according to the manufacturer’s instructions. Two additional recombinant proteins were purchased: biotinylated BA.1 S-6P (ACROBiosystems, #SPN-C82Ee) and Hu-1 S-2P (Miltenyi, #130-127-683).

### Flow cytometry and single-cell sorting

PBMCs were isolated from venous blood by standard density gradient centrifugation and used following cryopreservation at -150 °C. Cells were thawed in RPMI-1640 (Gibco)-10% FBS (Gibco), washed twice and incubated with a mixture of Hu-1, B.1.351, BA.1 S-6P and Hu-1 RBD tetramers in 100 µL of PBS (Gibco)-2% FBS during 40 min on ice. For cell sorting, cells were stained with 500 ng of Hu-1 spike BUV395-streptavidin, 500 ng of B.1.351 spike APC-streptavidin, 500 ng of BA.1 spike PE-streptavidin and 50 ng of Hu-1 RBD BV786-streptavidin. To exclude cells displaying nonspecific binding, a non-relevant tetramer was constructed using biotinylated bovine serum albumin coupled to BUV737-streptavidin. All tetramers were made just prior to the staining by incubating biotinylated proteins with fluorochrome-conjugated streptavidin at 4:1 molar ratio for 1h at 4 °C. 2.4 ng of free biotin was then added for 10 additional minutes before mixing of the tetramer. Cells were then washed and resuspended in the same conditions, prior to incubating with the fluorochrome-conjugated Ab cocktail at pre-titrated concentrations (1:100 for CD19, CD21, CD11c, CD71, CD38, CD3, CD14 and IgD, 1:50 for CD27) for 20 min at 4 °C. Viable cells were identified using a LIVE/DEAD Fixable Aqua Dead Cell Stain Kit (Thermo Fisher Scientific, 1:200) incubated with conjugated Abs. Samples were acquired using a LSR Fortessa SORP (BD Biosciences). For cell sorting, cells were stained using the same protocol and then sorted in 96-well plates using an Aria II cell sorter (BD Biosciences). Data were analyzed using FlowJo.

### UMAP

For UMAP generation and visualization (**Fig. S1a to d**), viable dump- CD19^+^ IgD^-^ cells from each sample included in the final analysis (**Table S2**) were first down-sampled to maximum 6,000 cells per sample. The UMAP (v3.1) plugin in FlowJO was then used on a concatenated FCS file containing all donors and time points to calculate the UMAP coordinates for the resulting 251,573 cells (with 15 nearest neighbors, metric = euclidean and minimum distance = 0.5 as default parameters), considering fluorescent intensities from the following parameters: FSC-A, SSC-A, CD19, CD21, CD11c, CD71, CD38 and CD27, while excluding the dump (CD3 and CD14), IgD, viability and tetramers channels. Based on surface markers expression, several clusters of pre-established B cell populations were manually gated separately: activated B cells (ABCs), resting MBCs, Double Negative (DN) 1, 2, 3 and 4, and antibody secreting cells (ASCs) (**Fig. S1f, g and h**). Contour plots (equal probability contouring, with intervals set to 5% of gated populations) of UMAP coordinates for each manually gated population (**Fig. S1c**) were further overlaid on the full UMAP projection in Adobe Illustrator (**Fig. S1d**). For visualization purposes, only the outermost density representing 95% of the total gated cells was kept for the final figure, all other contour lines were removed in Adobe Illustrator.

### Single-cell culture

Single-cell culture was performed as previously described(Crickx et al. 2021). Single B cells were sorted in 96-well plates containing MS40L^lo^ cells expressing CD40L (kind gift from G. Kelsoe(X. M. Luo et al. 2009)). Cells were co-cultured at 37 °C with 5% CO_2_ during 21 or 25 days in RPMI-1640 (Invitrogen, Thermo Fisher Scientific) supplemented with 10% HyClone FBS (Thermo Fisher Scientific), 55 µM 2-mercaptoethanol, 10 mM HEPES, 1 mM sodium pyruvate, 100 units/mL penicillin, 100 µg/mL streptomycin, and MEM non-essential amino acids (all from Invitrogen, Thermo Fisher Scientific), with the addition of recombinant human BAFF (10 ng/mL), IL-2 (50 ng/mL), IL-4 (10 ng/mL), and IL-21 (10 ng/mL; all from PeproTech). Part of the supernatant was carefully removed at days 4, 8, 12, 15 and 18 and the same amount of fresh medium with cytokines was added to the cultures. After 21 or 25 days of single-cell culture, supernatants were harvested and stored at -20 °C. Cell pellets were placed on ice and gently washed with PBS (Gibco, Thermo Fisher Scientific) before being resuspended in 50 µL of RLT buffer (QIAGEN) supplemented with 1% β-mercaptoethanol and subsequently stored at -80 °C until further processing.

### ELISA on single-cell culture supernatants

Total IgG and SARS-CoV-2 Hu-1 RBD, B.1.351 and BA.1 RBD specific IgG from culture supernatants were measured using homemade ELISA. 96-well ELISA plates (Thermo Fisher Scientific) were coated with either goat anti-human Ig (10 μg/mL, Invitrogen, Thermo Fisher Scientific) or recombinant SARS-CoV-2 Hu-1, B.1.351 or BA.1-RBD protein (2.5 µg/mL each) in sodium carbonate during 1 h at 37 °C. After plate blocking, cell culture supernatants were added for 1 h, then ELISA were developed using HRP-goat anti-human IgG (1 μg/mL, Immunotech) and TMB substrate (Eurobio). OD_450_ and OD_620_ were measured, and Ab-reactivity was calculated after subtraction of blank wells. Supernatants whose ratio of OD_450_-OD_620_ over control wells (consisting of supernatant from wells that contained spike-negative MBCs from the same single-cell culture assay) was over 10 were considered as positive for Hu-1 RBD, B.1.351 or BA.1 RBD. PBS was used to define background OD_450_-OD_620_.

### Single-cell IgH sequencing

Clones whose culture had proven successful (IgG concentration ≥ 1 µg/mL at day 21-25) were selected and RNA was extracted using the NucleoSpin96 RNA extraction kit (Macherey-Nagel) according to the manufacturer’s instructions. A reverse transcription step was then performed using the SuperScript IV enzyme (Thermo Fisher Scientific) in a 14 μL final volume (42 °C 10 min, 25 °C 10 min, 50 °C 60 min, 94 °C 5 min) with 4 µL of RNA and random hexamers (Thermo Fisher Scientific). A PCR was further performed based on the protocol established by Tiller *et al*.(Tiller et al. 2008). Briefly, 3.5 μL of cDNA was used as template and amplified in a total volume of 40 μL with a mix of forward L-V_H_, L-Vλ or L-Vκ primers and reverse Cγ, Cλ or Cκ primers for heavy chains, λ and κ light chains, respectively, using the HotStart^®^ Taq DNA polymerase (QIAGEN) and 50 cycles of PCR (94 °C 30 s, 58 °C 30 s, 72 °C 60 s). PCR products were sequenced with the reverse primer CHG-D1 and read on ABI PRISM 3130XL genetic analyzer (Applied Biosystems). Sequence quality was assessed with CodonCode Aligner software (CondonCode Corporation) or directly using the sangeranalyseR R package v1.14.0. 5’

#### PCR1 Primer Mix

**IgV_H_**

5ʹ L-VH 1 : ACAGGTGCCCACTCCCAGGTGCAG

5ʹ L-VH 3 : AAGGTGTCCAGTGTGARGTGCAG

5ʹ L-VH 4/6 : CCCAGATGGGTCCTGTCCCAGGTGCAG

5ʹ L-VH 5 : CAAGGAGTCTGTTCCGAGGTGCAG

5’L-VH 2 : CCTTCATGGGTCTTGTCCCAGATCACC

5’ L-VH 6 : CCATGGGGTGTCCTGTCACAGGTACAG

5’ L-VH 7 : GCAACAGGTGCCCACTCCCAGGTGCAG

**IgVκ**

5ʹ L-Vκ 1/2 : ATGAGGSTCCCYGCTCAGCTGCTGG

5ʹ L-Vκ 3 : CTCTTCCTCCTGCTACTCTGGCTCCCAG

5ʹ L-Vκ 4 : ATTTCTCTGTTGCTCTGGATCTCTG

**IgVλ**

5ʹ L-Vλ 1 : GGTCCTGGGCCCAGTCTGTGCTG

5ʹ L-Vλ 2 : GGTCCTGGGCCCAGTCTGCCCTG

5ʹ L-Vλ 3 : GCTCTGTGACCTCCTATGAGCTG

5ʹ L-Vλ 4/5 : GGTCTCTCTCSCAGCYTGTGCTG

5ʹ L-Vλ 6 : GTTCTTGGGCCAATTTTATGCTG

5ʹ L-Vλ 7 : GGTCCAATTCYCAGGCTGTGGTG

5ʹ L-Vλ 8 : GAGTGGATTCTCAGACTGTGGTG

3’ PCR1 Primers

**IgV_H_**

3ʹ Cγ (IgG) CH1 : GGAAGGTGTGCACGCCGCTGGTC

**IgVκ**

3ʹ Cκ 543 : GTTTCTCGTAGTCTGCTTTGCTCA

**IgVλ**

3ʹ Cλ : CACCAGTGTGGCCTTGTTGGCTTG

### Computational analyses of VDJ sequences

Processed FASTA sequences from the sequencing of heavy and light chains from sorted single-cells were annotated with Igblast v1.22.0 against the human ImMunoGeneTics (IMGT) reference database. Clonal cluster assignment (DefineClones.py) was performed with the Immcantation/Change-O toolkit on all heavy-chain V sequences. Sequences that had the same V_H_-gene, same J_H_-gene, including ambiguous assignments, and the same HCDR3 length with a maximal length-normalized nucleotide hamming distance of 0.15 were considered likely members of the same clonal group. Clonal groups were further corrected based on information from available light chains using the resolveLightChains() function from the Dowser v2.3 R package, prior to germline reconstruction using the createGermline() function from the same package. Mutation frequencies in V_H_ genes (**Fig. S2a**) and V_H_ gene distributions (**Fig. S2d**) were then calculated using the calcObservedMutations() and countGenes() functions from the SHazaM v1.2.0 R packages. Public datasets for naive and IgG^+^ cells from healthy donors were downloaded from the cAb-Rep database(Guo et al. 2019) and included donors from Vander Heiden, J. A. et al.(Vander Heiden et al. 2014), Rubelt, F. et al.(Rubelt et al. 2016), Galson, J. et al.(Galson et al. 2016) and Briney, B. et al.(Briney et al. 2019). Repertoire diversities statistics (**Fig. S2c**) were quantified by calculating Simpson diversities using the diversity() function from the Vegan v2.9-2 R package following rarefaction to equal sequence depth with repeated subsampling. Donut charts (**Fig. S2b**), generated with the circlize v0.4.16 R package, represent the clonal distribution of B cells from indicated samples/donors, with slices size corresponding to, and arranged according to, clones’ sizes. Total number of sequences is indicated in the middle of each donut chart, and slices are colored according to clone rank. The outer black semi-circle indicates the proportion of sequences belonging to expanded clones (two or more clonally related sequences), with frequencies further indicated on the top right of the donut plot. All analyses were performed with R v.4.4.1 on a macOS Sonoma 14.4.1 (Apple M1 Pro) system. Dedicated wrapper functions used for repertoire analysis are available here: https://github.com/PChappert/RERB.

### Gene synthesis, cloning, and mAb production

For HI-1 mAb (corresponding to CVB3_P1_G12 supernatant), IgV_H_ and V_L_ genes were synthesized in pUC19-Igγ1 or pUC-κ/pUC-λ expression (Synbio Technologies). HI-1 mAb was produced by transient transfection of V_H_-C_H_ and V_L_-C_L_ expression plasmids into exponentially growing Freestyle HEK 751 293-F, cultured in serum-free FreeStyle Expression Medium (Life Technologies) in suspension at 37 °C in a humidified 8 % CO_2_ incubator on a shaker platform rotating at 110 rpm. Twenty-four hours prior to transfection, cells were pelleted (300 xg, 5 min), resuspended at 1 x 10^6^ cells/mL in fresh expression medium, and cultured overnight under the same conditions. For mAb production, 50 μg of each V_H_ and V_L_ expressing plasmids were diluted in 100 µL of FectoPRO reagent (Polyplus) at a final DNA concentration of 0.8 μg/mL, followed by incubation for 10 min at room temperature before addition to the cell culture. At 24 h post-transfection, cells were diluted 1:1 with expression medium. Supernatants were harvested after 6 days, centrifuged at 1800 x g for 40 min, and filtered (0.2 μm). MAbs were purified by affinity chromatography using an AKTA pure FPLC instrument (GE Healthcare) on a HiTrap Protein A Column (GE Healthcare) and desalted 762 on a HiTrap desalting column (GE Healthcare).

LO-1 and LO-2 antibodies (corresponding to CVB4_P1_F7 and CVB3_P2_D3 supernatant respectively) were re-expressed by ProteoGenix SAS. Briefly, the cDNAs encoding the variable regions of the heavy and light chains were chemically synthesized, with optimization for expression in CHO cells, and subcloned in ProteoGenix’s proprietary mammalian cell expression vectors containing backbones for the human IgG1 heavy chain constant region and the human kappa/lambda light chain constant region. An endotoxin-free DNA preparation method was used for the construction of each vector and the vectors were then used to transfect XtenCHO cells with the XtenCHO transfection protocol, in a total volume of 3.5 mL. The culture medium was collected 8 days after transfection and purified by one-step affinity purification (Protein A). Final quality control was performed by spectrophotometric measurements of A_280nm_ and qualitative and quantitative analyses by SDS-PAGE.

### Reference antibodies

EY6A(D. Zhou et al. 2020) and S2H97(Starr et al. 2021) anti-RBD antibodies were obtained from ProteoGenix SAS. Purified parental research IgG1 versions of benchmarked anti-RBD antibodies (ADG2(Rappazzo et al. 2021), CB6(Shi et al. 2020), LY-CoV555(Jones et al. 2021), REGN10933 and REGN10987(Hansen et al. 2020)) were produced following cloning of synthetic DNA fragments encoding immunoglobulin variable domains (GeneArt; Thermo Fisher Scientific), as previously described(Planchais et al. 2022).

### Affinity measurement using biolayer interferometry (Octet)

#### Affinity measurement against RBDs

This high-throughput kinetic screening of supernatants using single antigen concentration has recently been extensively tested and demonstrated excellent correlation with multiple antigen concentration measurements.

Affinity for Hu-1, B.1.351 and BA.1 RBDs was assessed by biolayer interferometry (BLI) assays using an Octet Red96 instrument (Sartorius). Biosensors were equilibrated for 10 minutes in 1x PBS buffer with 0.1% BSA and 0.01% Tween 20 surfactant (PBS-BT) prior to measurement. Anti-Human Fc Capture (AHC) biosensors (18-5060) were immersed in supernatants from single-cell MBC cultures (or reference mAbs) at 25 °C for 500 s. Association was allowed to occur for 600 s in PBS-BT with Hu-1 or variant RBD (B.1.351, BA.1) at 100 nM, followed by dissociation for 900 s in PBS-BT. Between cycles, biosensors were regenerated via three successive alternating 30 s exposures to regeneration buffer (glycine HCl, 10 mM, pH 2.0) and PBS-BT (30 s each). Signals were double-referenced by subtracting both the reference sensor (unloaded sensor) and reference well (additional association cycle in the absence of ligand). Global curve fitting was then performed using a 1:1 binding model in HT Data analysis software v11.1 (ForteBio), to derive K_D_.

Sensors with response values (maximum RBD association) below 0.1 nM were considered non-binding. Hu-1 RBD non-binding mAbs (n = 2/124) were excluded from further analysis. For variant RBD non-binding mAbs, sensor-associated data (mAb loading and response) were manually checked to ensure that this was not the result of poor mAb loading. Monoclonal antibodies in culture supernatants were classified based on their dissociation constant (K_D_, M) as follows: high-binders (K_D_ < 10^-9^ M), mid-binders (10^-9^ ≤ K_D_ < 10^-8^ M), low-binders (10^-8^ ≤ K_D_ < 10^-7^ M), or non-binders (K_D_ ≥ 10^-7^ M). MAbs were defined as affected against a given variant RBD if the ratio of calculated K_D_ value against that RBD variant and the Hu-1 RBD was superior to three.

#### Affinity measurement against spike trimers

Following incubation at 37 °C (0 to 48 h kinetic), binding to Hu-1 or B.1.351 spike variants with either 2 or 6 proline substitutions was assessed by biolayer interferometry (BLI) using an Octet Red96 instrument (Sartorius). Biosensors were equilibrated for 10 minutes in 1x PBS buffer with 0.1% BSA and 0.01% Tween 20 surfactant (PBS-BT) prior to measurement at 37 °C. Anti-Human Fc Capture (AHC) biosensors (18-5060) were immersed in supernatants from single-cell MBC cultures (or reference mAbs) at 37 °C for 500 s. Association was allowed to occur for 600 s in PBS-BT with Hu-1 or variant spike trimers (B.1.351, BA.1) at 33 nM, followed by dissociation for 1200 s in PBS-BT. Between cycles, biosensors were regenerated via three successive alternating 30 s exposures to regeneration buffer (glycine HCl, 10 mM, pH 2.0) and PBS-BT (30 s each). Signals were double-referenced by subtracting both the reference sensor (unloaded sensor) and reference well (additional association cycle in the absence of ligand). Global curve fitting was then performed using a 1:1 binding model in HT Data analysis software v. 11.1 (ForteBio), to derive K_D,_ k_on_ and k_off_. For apparent K_D_ estimation global curve fitting was performed using the exact concentration of each trimer used. Trimer accessibility for a given mAb (**Fig. 5f**) was calculated by varying the concentration of each trimer used for global curve fitting until reaching a derived k_on_ matching the true k_on_ measured against the isolated RBD monomer of the same variant. Trimer accessibility is reported as a percent-converted ratio of corrected concentration (likely seen by the mAb) over exact concentration in the well. The percentage of accessible RBD epitope was calculated on three independent experiments.

### Affinity measurement using Surface Plasmon Resonance (SPR)

#### Affinity measurement against spike trimers

CM5 chips (Cytiva) were primed using PBS, washed 3 times with NaOH (50 mM) and SDS (0.1%) 180 s each time (5 µL/min). At least 10,000 RU of anti-human IgG Fc capture antibody (Cytiva) was covalently bound to CM5 chip by activating using 1:1 NHS:EDC (600 s, (5 µL/min), injecting the capture antibody (900 s) in acetate buffer pH 5.5 and quenching with EtNH2 (1200 s, 5 µL/min). Then at least 100 RU of Covid-specific mAb (10 µg/mL) / or supernatant (diluted X100) from cell culture were injected on the first three canals and a non-specific IgG on the last canal as reference. The antigen (Hu-1 or B.1.351 S-6P) was injected at different concentrations (30 µL/min). Between each injection the canals were regenerated by three injections of MgCl_2_ (3 M, 120 s). Signal was subtracted to the reference canal bound with a non-specific antibody. Experiment was done at 25 °C. K_D,_ k_on_ and k_off_ were determined using a kinetic analysis (Langmuir binding model) with Biacore T200 evaluation software.

### Epitope-binning assays

Epitope-binning classical sandwich assays (**Fig. 3a and S3a**) were performed using biolayer interferometry (BLI) on an Octet Red96 instrument (Sartorius). 94 MBC culture supernatants (loaded first) were tested against a set of four reference mAbs covering the RBM: class 1 CB6(Shi et al. 2020); class 2 LY-CoV555(Jones et al. 2021); class 3 REGN10987(Hansen et al. 2020); and class 1/4 ADG2(Rappazzo et al. 2021; Yuan et al. 2022) for binding to the Hu-1 RBD (**Fig. S3b**). As control, three re-expressed mAbs (class 2/3 HI-1, class 4 LO-1 and class 5 LO-2) and 7 reference mAbs (CB6, LY-CoV555, REGN10987, ADG2, class 1 REGN10933(Hansen et al. 2020), class 4 EY6A(D. Zhou et al. 2020), and class 5 S2H97(Starr et al. 2021)) were included.

For each tested mAb, anti-human Fc capture (AHC) biosensors (18-5060) were first equilibrated for 10 minutes in 1x PBS buffer with 0.1% BSA and 0.01% Tween 20 surfactant (PBS-BT) prior to measurement at 30 °C. Next, biosensors were immersed in culture supernatants containing mAbs or in solution of re-expressed mAbs (5 µg/mL) at 30 °C for 600 seconds. Biosensors were then quenched by incubating loaded biosensors in polyclonal human IgG solution (10 µg/mL) for 300 s. Association with Hu-1 RBD was allowed to occur for an additional 300 s in PBS-BT at 100 nM. Competitive binding was then assessed against one of the predefined set of four reference mAbs covering the RBM as described above (5 µg/mL) at 30 °C for 300 s. Biosensors were then regenerated by three rounds of alternating cycles of regeneration buffer (10 mM glycine HCl, pH 2.0) for 30 s and PBS-BT for 30 s. 3 additional cycles of mAb loading, human IgG quenching, RBD binding, competitive binding and regeneration steps were further performed on the same biosensor to test competitive binding to all four references in row for a given mAb. Heatmap visualization of epitope-binning results for culture supernatants were generated using the ComplexHeatmap v. 2.20.0 R package, with additional unsupervised row clustering (**Fig. 3a**).

Reciprocal epitope-binning assays (**Fig. S3c**) was performed in the same manner on all re-expressed mAbs with all reference mAbs, and heatmap visualization was generated using pheatmap: Pretty Heatmaps v. 1.0.12.

### Structural prediction of mAb/Hu-1 RBD complexes

All structure predictions were performed using an in house implementation of the AlphaFold3 algorithm(Abramson et al. 2024), entering the full heavy chain and the full light chain VDJ amino-acid sequences (beginning of FR1 to end of FR4 with (Fab) or without (Fv) the IGKC, IGLC2 or IGHG1 C_H1_ domains, with reversion to germline when needed for missing early or end parts of the original FR1 or FR4 sequences due to low sequencing quality at the site of primer hybridization) and the amino-acid sequence of Hu-1 RBD, previously used for all epitope-binning experiments.

**RBD(Hu1):** RVQPTESIVRFPNITNLCPFGEVFNATRFASVYAWNRKRISNCVADYSVLYNSASFSTFKCYGVSPTKLNDLCFTNVYADSFVIRGDEVRQIAPGQTGKIADYNYKLPDDFTGCVIAWNSNNLDSKVGGNYNYLYRLFRKSNLKPFERDISTEIYQAGSTPCNGVEGFNCYFPLQSYGFQPTNGVGYQPYRVVVLSFELLHAPATVCGPKKSTNLVKNKCVNF;

**IGHG1(CH1_domain):** ASTKGPSVFPLAPSSKSTSGGTAALGCLVKDYFPEPVTVSWNSGALTSGVHTFPAVLQSSGLYSLSSVVTVPSSSLGTQTYICNVNHKPSNTKVDKKV;

**IGKC:** RTVAAPSVFIFPPSDEQLKSGTASVVCLLNNFYPREAKVQWKVDNALQSGNSQESVTEQDSKDSTYSLSSTLTLSKADYEKHKVYACEVTHQGLSSPVTKSFNRGEC;

**IGLC2:** GQPKAAPSVTLFPPSSEELQANKATLVCLISDFYPGAVTVAWKADSSPVKAGVETTTPSKQSNNKYAASSYLSLTPEQWKSHRSYSCQVTHEGSTVEKTVAPTECS.

The structure template option was only used for the RBD and a single random seed was used for all predictions. All data and structures reported in this study correspond to model 0 of the five ranked predictions provided in the output of the AlphaFold3 server. Based on the AlphaFold3 server recommendations, confidence score distribution in our dataset (**Fig. S4b**) and prior experience(Fournier et al. 2026), the AlphaFold3-predicted structures were classified as follows: predictions for which both RBD/heavy chain and RBD/light chain “chain_pair_ipTM” confidence scores ≥ 0.8 were considered to be of “very high confidence”, predictions with both RBD/heavy chain and RBD/light chain “chain_pair_ipTM” confidence scores ≥ 0.6, but at least one < 0.8, were considered to be “confident predictions” and the confidence level for all other predictions was considered to be “poor” and the latter were discarded. These predictions were done in parallel for both the Fab and the Fv form of each IgVH and IgVl pairs, only “very high confidence” and “confident” predictions were retained for further analysis (**Fig. S4b**). They had global ipTM confidence scores > 0.74 for Fvs and > 0.75 for Fabs (**Table S4b**). Confidence scores for all predictions are shown in **Table S4a**. One representative antibody per class was selected to illustrate the predicted structures (**Fig. S4a**), with coloring based on per-atom confidence as estimated by the predicted local distance difference test (pLDDT) confidence score.

For epitope mapping (**Fig. S4e and Table S4b**), Fv-based predictions were used when available, as they displayed slightly higher ipTM confidence scores overall (**Fig. S4b**), but a few Fab-based predictions were also included for pairs with “poor” Fv-based predictions. For intra-spike clashes and bivalency predictions (**Fig. 4, S5 and Table S4**), only Fab-based predictions were used.

Representations for all predicted structures were created within UCSF ChimeraX v.1.10(Meng et al. 2023), developed by the Resource for Biocomputing, Visualization, and informatics at the University of California, San Francisco, with support from National Institutes of Health R01-GM129325 and the Office of Cyber Infrastructure and Computational Biology, National Institute of Allergy and Infectious Diseases.

Targeted residues heatmap (**Fig. S4e)** report the maximum predicted probability of individual residues from heavy or light chains being in contact with the indicated residue in the RBD protein (8Å between the representative atom for each residue(Abramson et al. 2024); AlphaFold3 “contact_probs” full array output). Raw contact probabilities for the heavy chain and light chain respectively are reported in **Table S4b**. Unsupervised hierarchical clustering of all tested mAbs was performed with the maximal predicted contact probabilities for the heavy and light chains and a k value of 7 for the dendextend package’s cutree() function. AlphaFold3-predicted structures for antibodies described in previous publications are also reported (ADG2 (ADG20, PDB 7U2D)(Rappazzo et al. 2021; Yuan et al. 2022), REGN10933 (PDB 6XDG)(Hansen et al. 2020), EY6A (PDB 6ZER)(D. Zhou et al. 2020), and S2H97 (PDB 7M7W)(Starr et al. 2021) mAbs).

All AlphaFold3 downstream analyses were performed with R v.4.3.1, using the following packages and libraries: rjson v.0.2.21, ggplot2 v.3.5.1, dendextend v.1.17.1, ComplexHeatmap v.2.18.0. Dedicated wrapper functions used for interaction with AlphaFold3 models are available here: https://github.com/PChappert/RERB.

#### Anti-RBD Fab-spike interactions modeling

For intra-Fab distances and intra-spike clashes (**Fig. 5**), confident Fab/Hu-1 RBD models predicted by AlphaFold3 (n = 49) were first aligned to the three RBDs of four experimentally resolved (cryo-EM) SARS-CoV-2 spike trimer structures using the Matchmaker command implemented in UCSF ChimeraX v.1.10(Meng et al. 2023). Used structures were comprised of canonical spike trimers with zero (PDB 7BNM), one (PDB 7BNN) or two (PDB 7BNO) RBDs in the “up” conformation(Benton et al. 2021) and an expanded open trimer stabilized upon antibody binding (S304 mAb, PDB 7JW0(Piccoli et al. 2020)). Intra-spike steric clashes for each RBD-bound Fab were computed using the clashes sel save() function in UCSF ChimeraX command-line interface. To solely focus on steric clashes with residues outside of the bound RBD, residues 333-526 from the bound spike monomer were excluded from the analysis. Each mAb was successively analyzed bound to all three RBDs of each spike, and binding was considered possible for this mAb to that spike trimer conformation if binding to at least one of the three RBDs was predicted to occur without any measured clash.

To assess whether bivalent engagement of mAbs on any of these four prefusion spike trimer conformations was structurally feasible, we further selected all mAbs for whom two Fabs were predicted to bind simultaneously on individual RBDs without any measured steric clash. We then measured the distances between the Cα atoms of the last lysine residues in both C_H1_ domains, near the C-termini of adjacent Fab C_H1_ domains using the distance() function through the command-line interface in UCSF ChimeraX, v.1.10(Meng et al. 2023). We further used a maximum 65Å distance cutoff to identify mAbs with bivalency potential, as previously defined for other anti-RBD IgG mAbs by Barnes *et al*. based on analogous distances in crystal structures of intact IgGs(Barnes et al. 2020).

### Illustration tool

All figures were assembled using Adobe Illustrator. Fig. 1a (Broutin, M. (2026) https://BioRender.com/j2zaod4) and 2a (Broutin, M. (2026) https://BioRender.com/xt4sk77) were created using BioRender.

### Statistical analyses

Mann-Whitney test, repeated measures mixed effects model analysis (with Tukey’s correction), Kruskal-Wallis test (with Dunn’s correction), simple linear regression, Chi-square test and individual Fisher’s test with Bonferroni correction were used to compare continuous variables as appropriate (indicated in figures). A *p*-value ≤ 0.05 was considered statistically significant. Statistical analyses were all performed using GraphPad Prism 11 (La Jolla, CA, USA).

## ACKNOWLEDGMENTS

The authors are grateful to the personal of the technological core facilities of the SFR Necker (InsermUS24/CNRS UAR3633) including the cytometry platform for help on single-cell sorting and Chiara Guerrera from the proteomics platform for providing access to an OCTET Red96 instrument. The authors would like to thank the staff of the University Clinical Research Platform for Eastern Paris (URCEST, CRCEST, CRB AP-HP. SU, Saint-Antoine Hospital, AP-HP), and would particularly like to acknowledge the contribution of Amel Touati, Alexandra Rousseau, Nacilla Haicheur and Lucie Armenoult.

## AUTHOR CONTRIBUTIONS

Conceptualization: M.B., A.S., M.V., O.L., J.C.W., P.C., M.M.

Methodology: M.B., A.S., M.V., P.B., F.A.R., H.M., O.S., P.C., M.M.

Investigation: M.B., A.S., I.A., A.D., D.P., M.V., A.A., I.F., A.V., M.C., J.M.

Visualization: M.B., A.S., P.C., M.M.

Funding acquisition: O.L., P.C., M.M.

Project administration: P.C., M.M.

Supervision: J.C.W., P.C., M.M.

Writing – original draft: M.B., A.S., P.C., M.M.

Writing – review & editing: All the authors

