## Supplemental Figures for "Variant-specific spike conformational dynamics shape memory B cell selection during recall"

### Fig. S1

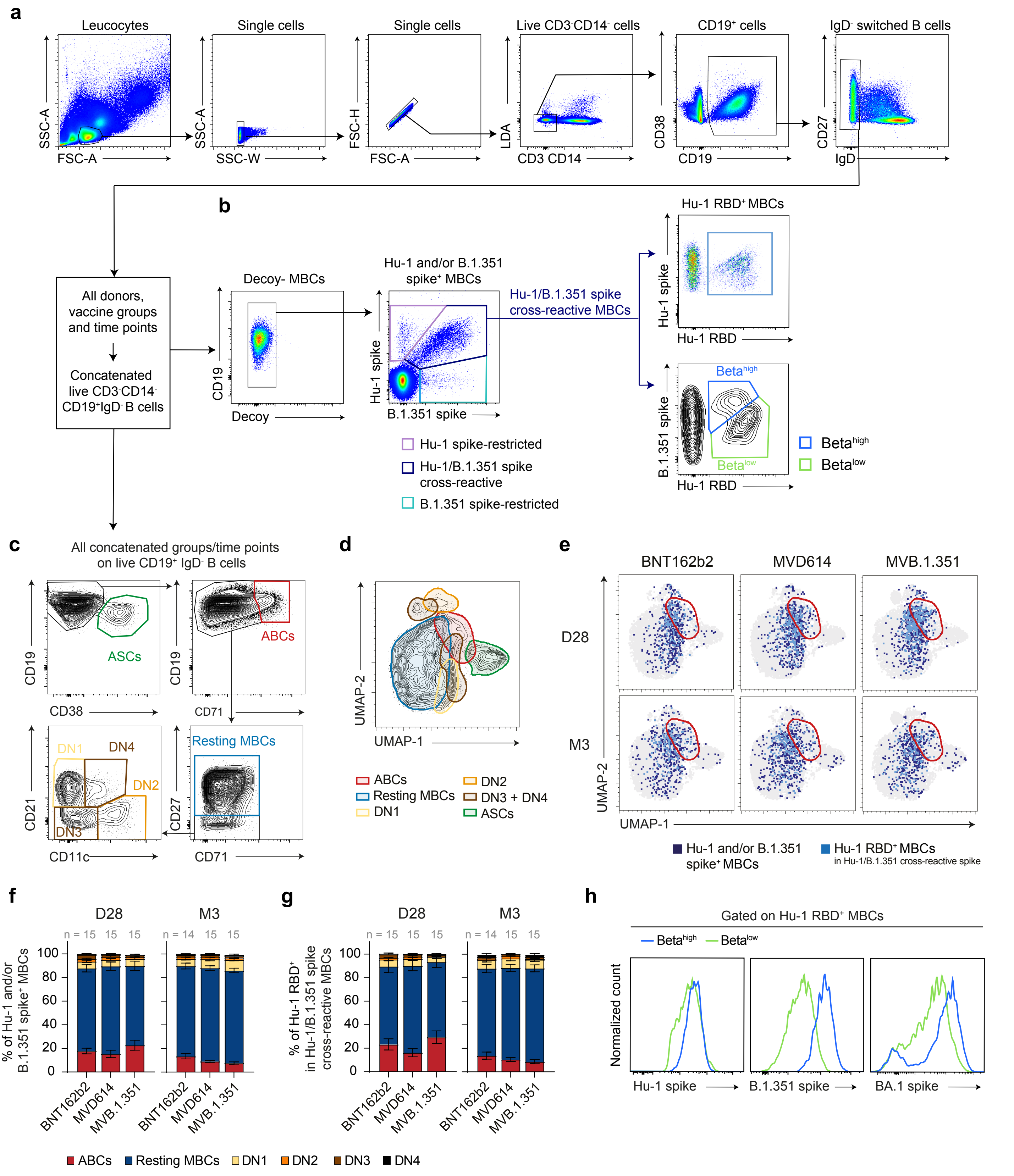

a
Leucocytes
Single cells
Single cells
Live CD3-CD14- cells
CD19+ cells
IgD- switched B cells
SSC-A
FSC-A
SSC-W
FSC-H
LDA
CD3 CD14
CD38
CD19
CD27
IgD
b
All donors, vaccine groups and time points
Concatenated live CD3-CD14- CD19+IgD- B cells
Decoy- MBCs
CD19
Decoy
Hu-1 and/or B.1.351 spike+ MBCs
Hu-1 spike
B.1.351 spike
Hu-1/B.1.351 spike cross-reactive MBCs
Hu-1 spike-restricted
Hu-1/B.1.351 spike cross-reactive
B.1.351 spike-restricted
Hu-1 RBD+ MBCs
Hu-1 RBD
Beta high
Beta low
c
All concatenated groups/time points on live CD19+ IgD- B cells
CD19
CD38
ASCs
CD71
ABCs
CD21
CD11c
DN1
DN4
DN2
DN3
CD27
Resting MBCs
d
UMAP-2
UMAP-1
ABCs
Resting MBCs
DN1
DN2
DN3 + DN4
ASCs
e
BNT162b2
MVD614
MVB.1.351
D28
M3
Hu-1 and/or B.1.351 spike+ MBCs
Hu-1 RBD+ MBCs in Hu-1/B.1.351 cross-reactive spike
f
D28
M3
n = 15 15 15
n = 14 15 15
% of Hu-1 and/or B.1.351 spike+ MBCs
0
20
40
60
80
100
BNT162b2
MVD614
MVB.1.351
g
% of Hu-1 RBD+ in Hu-1/B.1.351 spike cross-reactive MBCs
ABCs
Resting MBCs
DN1
DN2
DN3
DN4
h
Gated on Hu-1 RBD+ MBCs
Beta high
Beta low
Normalized count
Hu-1 spike
B.1.351 spike
BA.1 spike

**Fig. S1 (relates to Fig. 1). Flow cytometric gating and phenotypic analysis of RBD- and spike-specific B cell subsets in the COVIBOOST cohort.**

(**a**) Flow cytometry gating strategy for donors' PBMCs analysis. After doublet exclusion, CD19$^+$ B cells were gated within live CD3$^-$CD14$^-$ cells and switched memory B cells were further gated as CD19$^+$IgD$^-$ cells (a). IgD$^-$CD19$^+$ switched mature B cells from samples from all donors, vaccine groups and time points included in the analysis were then concatenated for further analysis. **(b)** Within concatenated live CD19$^+$IgD$^-$ B cells, and after exclusion of decoy tetramer$^+$ B cells (left panel), Hu-1 or B.1.351 spike tetramer-binding B cells were further classified as Hu-1 spike-restricted (pastel purple), Hu-1/B.1.351 spike-cross-reactive (dark blue) or B.1.351 spike-restricted (cyan) (middle panel). Hu-1 RBD-specific cells (upper right panel) and Beta$^{high}$ (blue) and Beta$^{low}$ (green) subpopulations (lower right panel) were further gated among Hu-1/B.1.351 spike-cross-reactive cells, based on Hu-1 RBD and Hu-1 or B.1.351 spike tetramer stainings, respectively. (**c**) Within concatenated live CD19$^+$IgD$^-$ B cells, antibody secreting cells (ASCs) were defined as CD19$^{+/int}$ CD38$^{hi}$ B cells (upper left panel). Among non-ASCs, CD71$^{hi}$ B cells were defined as Activated B Cells (ABCs) (upper right panel). CD71$^{low/int}$ B cells were further subdivided into CD27$^+$ resting MBCs and CD27$^-$ Double Negatives (DNs) B cells (lower right panel), with the DN compartment resolved into CD11c$^-$CD21$^+$ (DN1), CD11c$^+$CD21$^-$ (DN2), CD11c$^-$CD21$^-$ (DN3), and CD11c$^+$CD21$^+$ (DN4) subsets (lower left panel). (**d**) Uniform Manifold Approximation and Projection (UMAP) of concatenated CD19$^+$IgD$^-$ B cells from samples of all donors, vaccine groups and time points included in the analysis. Position (95% contour plot level) of indicated populations is overlaid on the UMAP: ABCs (red), resting MBCs (blue), DN1 (yellow), DN2 (orange), DN3 and DN4 (brown), ASCs (green). (**e**) UMAPs of concatenated CD19$^+$IgD$^-$ B cells from samples of all donors separated by vaccine group and time point post-booster-vaccination, with all Hu-1 and/or B.1.351 spike-specific MBCs and Hu-1 RBD-specific Hu-1/B.1.351 spike-cross-reactive B cells overlaid in dark blue and light blue dots, respectively. The position (95% contour plot level) of $CD71^+$ ABCs is further outlined in red on the UMAP. (**f**) Hu-1 and/or B.1.351 spike-specific MBCs (left) and Hu-1 RBD-specific Hu-1/B.1.351 spike-cross-reactive MBCs (right) among the main B cells clusters defined in panel C, stratified by vaccine group and time point post-booster-vaccination. (**g**) Fluorescence intensity distributions of Hu-1, B.1.351 and BA.1 spike tetramer stainings on $Beta^{high}$ and $Beta^{low}$ MBCs.

Fig. S2

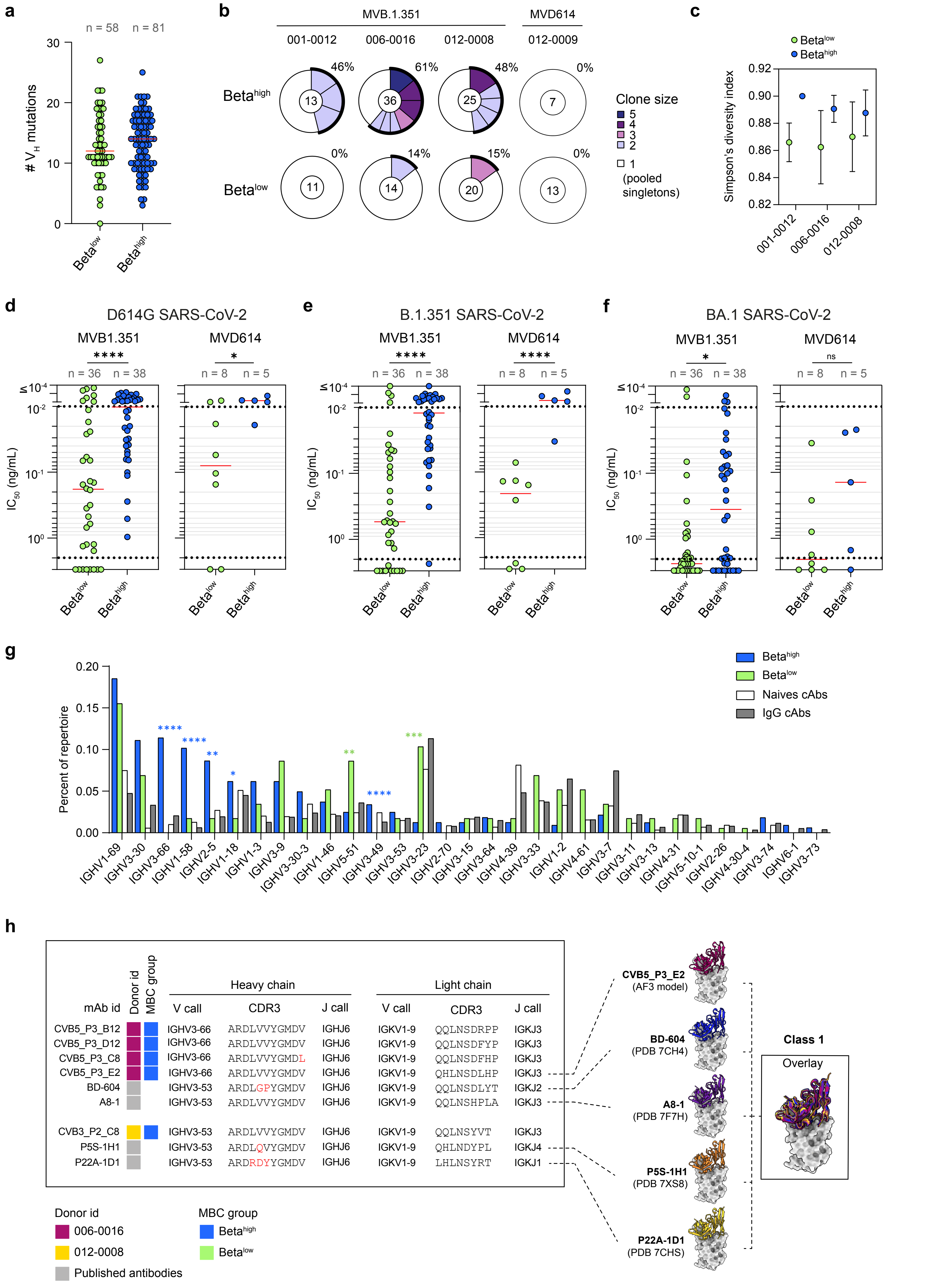

a
n = 58
n = 81
### V_H mutations
Beta^low
Beta^high
b
MVB.1.351
MVD614
001-0012
006-0016
012-0008
012-0009
Beta^high
Beta^low
46%
61%
48%
0%
14%
15%
Clone size
5
4
3
2
1 (pooled singletons)
c
Simpson's diversity index
001-0012
006-0016
012-0008
d
D614G SARS-CoV-2
e
B.1.351 SARS-CoV-2
f
BA.1 SARS-CoV-2
MVB1.351
MVD614
n = 36
n = 38
n = 8
n = 5
ns
IC50 (ng/mL)
g
Percent of repertoire
Beta^high
Beta^low
Naives cAbs
IgG cAbs
IGHV1-69
IGHV3-30
IGHV3-66
IGHV1-58
IGHV2-5
IGHV1-18
IGHV1-3
IGHV3-9
IGHV3-30-3
IGHV1-46
IGHV5-51
IGHV3-49
IGHV3-53
IGHV3-23
IGHV2-70
IGHV3-15
IGHV3-64
IGHV4-39
IGHV3-33
IGHV1-2
IGHV4-61
IGHV3-7
IGHV3-11
IGHV3-13
IGHV4-31
IGHV5-10-1
IGHV2-26
IGHV4-30-4
IGHV3-74
IGHV6-1
IGHV3-73
h
mAb id
Donor id
MBC group
Heavy chain
Light chain
V call
CDR3
J call
CVB5_P3_B12 IGHV3-66 ARDLVVYGMDV IGHJ6 IGKV1-9 QQLNSDRPP IGKJ3
CVB5_P3_D12 IGHV3-66 ARDLVVYGMDV IGHJ6 IGKV1-9 QQLNSDFYP IGKJ3
CVB5_P3_C8 IGHV3-66 ARDLVVYGMDL IGHJ6 IGKV1-9 QQLNSDFHP IGKJ3
CVB5_P3_E2 IGHV3-66 ARDLVVYGMDV IGHJ6 IGKV1-9 QHLNSDLHP IGKJ3
BD-604 IGHV3-53 ARDLGPYGMDV IGHJ6 IGKV1-9 QQLNSDLYT IGKJ2
A8-1 IGHV3-53 ARDLVVYGMDV IGHJ6 IGKV1-9 QQLNSHPLA IGKJ3
CVB3_P2_C8 IGHV3-53 ARDLVVYGMDV IGHJ6 IGKV1-9 QQLNSYVT IGKJ3
P5S-1H1 IGHV3-53 ARDLQVYGMDV IGHJ6 IGKV1-9 QHLNDYPL IGKJ4
P22A-1D1 IGHV3-53 ARDRDYYGMDV IGHJ6 IGKV1-9 LHLNSYRT IGKJ1
CVB5_P3_E2 (AF3 model)
BD-604 (PDB 7CH4)
A8-1 (PDB 7F7H)
P5S-1H1 (PDB 7XS8)
P22A-1D1 (PDB 7CHS)
Class 1
Overlay
Donor id
006-0016
012-0008
Published antibodies
MBC group
Beta^high
Beta^low

**Fig. S2 (relates to Fig. 2). Repertoire and neutralization potency of Beta$^{high}$ and Beta$^{low}$ MBCs.** (**a-c**) V(D)J repertoire analysis of single-cell sorted Beta$^{low}$ (green, n = 58) and Beta$^{high}$ (blue, n = 81) MBC populations from 4 donors (n = 1 and n = 3 from the MVD614 and MVB.1.351 vaccine groups, respectively). (**a**) Absolute number of mutations in the IgV$_H$ gene of MBCs from indicated populations pooled from all donors. Bars indicate median. (**b**) Pie chart showing the clonal distribution of Beta$^{high}$ and Beta$^{low}$ MBCs for each of the 4 donors. Colored slices correspond to clones of size ≥ 1, with area and color of the slice linked to clone size (number of unique cells). All singletons are grouped as one final white slice. The number at the center indicates the total number of sequenced cells per MBC population and donor. (**c**) Simpson's diversity index of Beta$^{high}$ and Beta$^{low}$ MBC populations for the 3 patients with at least 10 cells sequenced for both populations. Mean ± SD. (**d-f**) *In vitro* neutralization half-maximal inhibitory concentration (IC$_{50}$) and potency (high: IC$_{50}$ < 0.01 ng/mL; mid: ≥ 0.01 ng/mL to < 2 ng/mL; low/none ≥ 2 ng/mL) of Beta$^{high}$ and Beta$^{low}$ MBC-derived mAbs against authentic D614G (left), B.1.351 (middle) and BA.1 SARS-CoV-2 strains (right). mAbs are grouped based on donor's vaccine group. (**g**) Distribution of IgV$_H$ gene usage among Beta$^{high}$ (blue) and Beta$^{low}$ MBCs (green) from the 4 donors analyzed in (a-c), relative to naive (white) and IgG$^+$ B cells (grey) from public datasets. (**h**) Public class 1 RBD-specific IGHV3-53/66/IGKV1-9 clonotype identified in Beta$^{high}$ MBCs from two donors. The table (left) summarizes heavy and light chain V and J gene usage and CDR3 amino-acid sequences from monoclonal antibodies belonging to this clonotype isolated from indicated donor in our cohort or published datasets with linked PDB structures: BD-604 (PDB 7CH4; A8-1 (PDB 7F7H); P5S-1H1 (PDB 7XS8); P22A-1D1 (PDB 7CHS). Available PDB structures of antibody Fv domains bound to the Hu-1 RBD are displayed on the right of the table alongside the confident AlphaFold 3-

predicted complex for one the Beta$^{high}$ MBC-derived mAb (CVB5_P3_E2), with structural superposition of all complexes further shown on the far right.

(d-f) Mann-Whitney test ****$p < 0.0001$, *$p < 0.05$, ns = non-significant.

Fig. S3

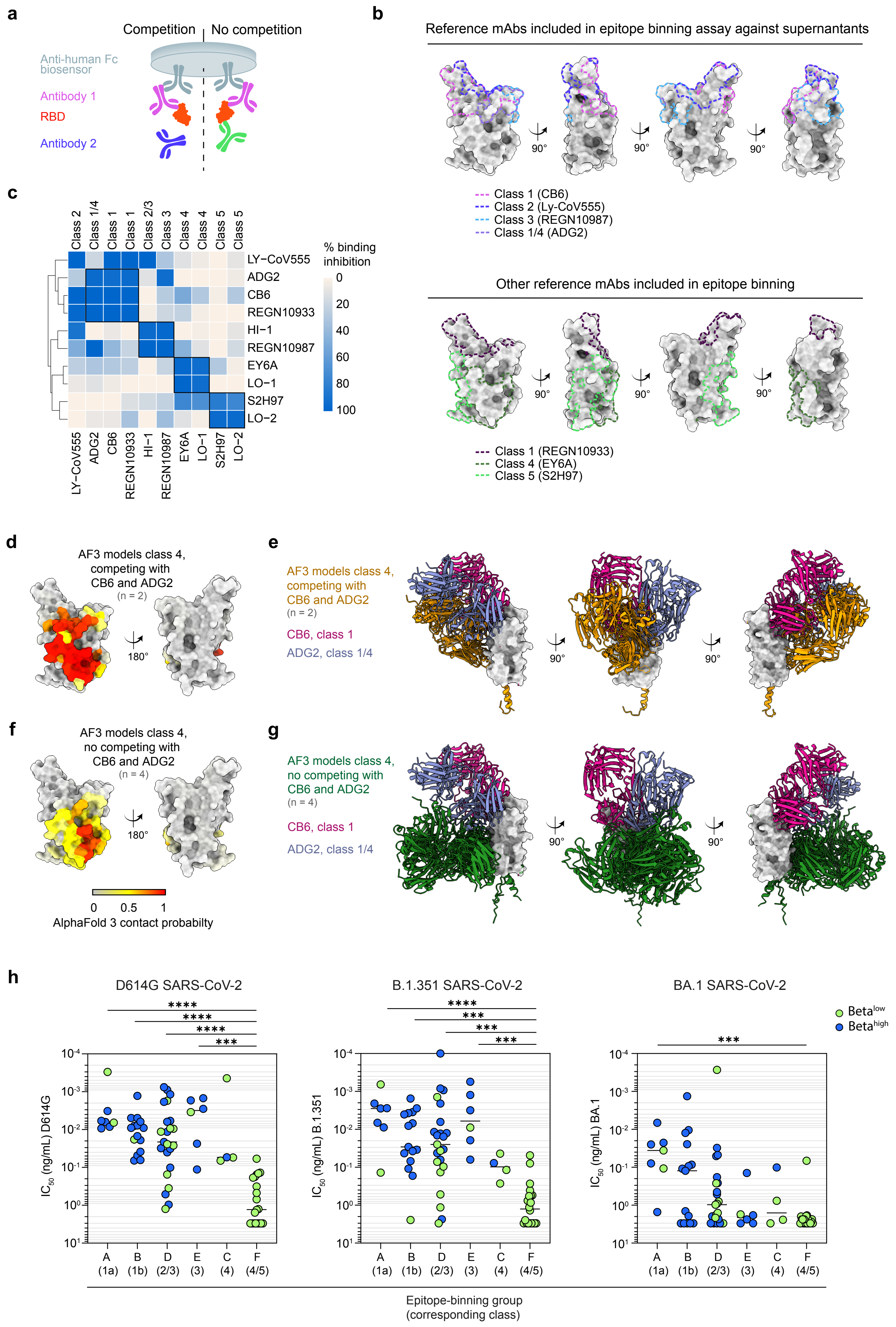

a
Competition
No competition
Anti-human Fc biosensor
Antibody 1
RBD
Antibody 2
b
Reference mAbs included in epitope binning assay against supernantants
90°
Class 1 (CB6)
Class 2 (Ly-CoV555)
Class 3 (REGN10987)
Class 1/4 (ADG2)
Other reference mAbs included in epitope binning
Class 1 (REGN10933)
Class 4 (EY6A)
Class 5 (S2H97)
c
Class 2
Class 1/4
Class 1
Class 1
Class 2/3
Class 3
Class 4
Class 4
Class 5
Class 5
LY−CoV555
ADG2
CB6
REGN10933
HI−1
REGN10987
EY6A
LO−1
S2H97
LO−2
% binding inhibition
0
20
40
60
80
100
d
AF3 models class 4, competing with CB6 and ADG2
(n = 2)
180°
e
AF3 models class 4, competing with CB6 and ADG2
(n = 2)
CB6, class 1
ADG2, class 1/4
f
AF3 models class 4, no competing with CB6 and ADG2
(n = 4)
0
0.5
1
AlphaFold 3 contact probabilty
g
AF3 models class 4, no competing with CB6 and ADG2
(n = 4)
CB6, class 1
ADG2, class 1/4
h
D614G SARS-CoV-2
B.1.351 SARS-CoV-2
BA.1 SARS-CoV-2
****
***
Beta^low
Beta^high
IC50 (ng/mL) D614G
IC50 (ng/mL) B.1.351
IC50 (ng/mL) BA.1
10^-4
10^-3
10^-2
10^-1
10^0
10^1
A (1a)
B (1b)
D (2/3)
E (3)
C (4)
F (4/5)
Epitope-binning group (corresponding class)

**Fig. S3 (relates to Fig. 3). Epitope definition of Beta$^{high}$- and Beta$^{low}$-derived mAbs by epitope-binning.**

(**a**) Epitope-binning strategy using a competitive binding (BLI) sandwich assay with Hu-1 RBD and tested mAbs loaded first on the biosensor. (**b**) Published epitopes for 7 anti-RBD class 1, 2, 3, 1/4, 4 and 5 reference mAbs (CB6: PDB 7C01; Ly-CoV555: PDB 7KMG; REGN10987 and REGN10933: PDB 6XDG; ADG20: PDB 7U2D; EY6A: PDB 6ZER; and S2H97: PDB 7M7W) used in the assay are outlined in colored dotted lines on top of Hu-1 RBD structure (PDB 6M0J). (**c**) Reciprocal competitive binding heatmap to Hu-1 RBD for 10 re-expressed anti-RBD mAbs. This included the 7 published mAbs described above and three re-expressed mAbs from our dataset: HI-1 (AlphaFold 3-predicted class 2/3 Beta$^{high}$-MBC-derived CVB3_P1_G12) and LO-1 and LO-2 (AlphaFold 3-predicted class 4 and 5a Beta$^{low}$-MBC-derived CVB4_P1_F7 and CVB3_P2_D3, respectively). Data from epitope-binning assay are expressed as percentage of binding inhibition: white denotes no inhibition, whereas dark-blue denotes complete inhibition. MAbs are hierarchically clustered based on their binding profiles (dendrogram). (**d-e)** Average AlphaFold 3-predicted contact probabilities projected on the Hu-1 RBD structure for predicted class 4 mAbs competing (d, n = 2) or not competing (e, n = 4) with class 1 CB6 and class 1/4 ADG2 reference mAbs in our epitope binning assay. (**f-g**) Overlayed AlphaFold 3-predicted structures of Hu-1 RBD, class 4 mAbs competing (f, n = 2, orange) or not competing (g, n = 4, green) with class 1 CB6 and class 1/4 ADG2 reference mAbs in our epitope binning assay and corresponding reference mAbs structures (CB6, purple and ADG2, blue). (**h**) *In vitro* neutralization half-maximal inhibitory concentration ($IC_{50}$) of Beta$^{high}$ (green) and Beta$^{low}$ (blue) MBC-derived mAbs against authentic D614G (left), B.1.351 (middle) and BA.1 SARS-CoV-2 strains (right). mAbs are grouped based on epitope binning group and colored according to the MBC population of origin. (h) Kruskal-Wallis test with Dunn's multiple-comparison correction. ****$p < 0.0001$, ***$p < 0.001$.

### Fig. S4

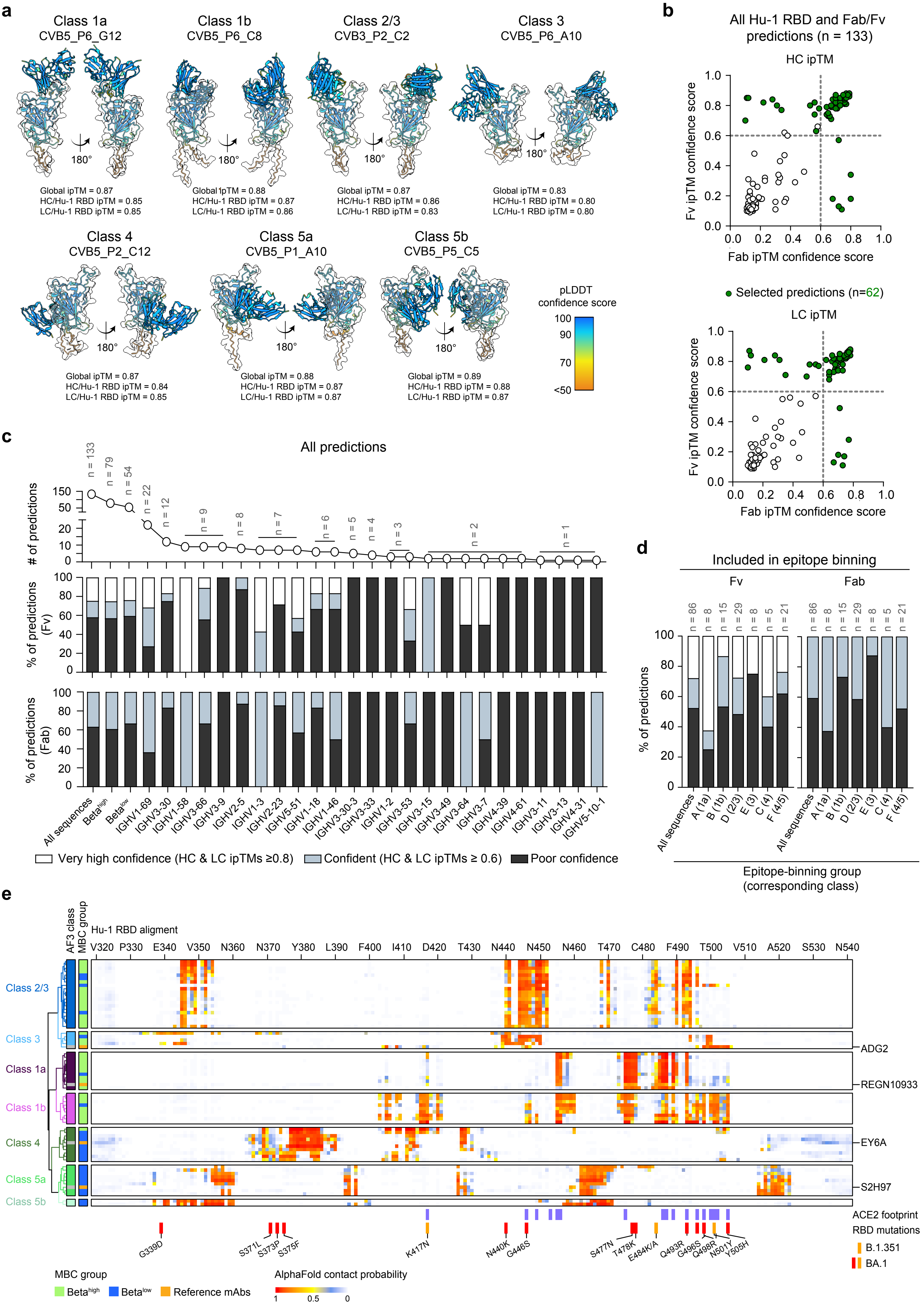

a
Class 1a CVB5_P6_G12
Class 1b CVB5_P6_C8
Class 2/3 CVB3_P2_C2
Class 3 CVB5_P6_A10
180°
Global ipTM = 0.87
HC/Hu-1 RBD ipTM = 0.85
LC/Hu-1 RBD ipTM = 0.85
Global ipTM = 0.88
HC/Hu-1 RBD ipTM = 0.87
LC/Hu-1 RBD ipTM = 0.86
Global ipTM = 0.87
HC/Hu-1 RBD ipTM = 0.86
LC/Hu-1 RBD ipTM = 0.83
Global ipTM = 0.83
HC/Hu-1 RBD ipTM = 0.80
LC/Hu-1 RBD ipTM = 0.80
Class 4 CVB5_P2_C12
Class 5a CVB5_P1_A10
Class 5b CVB5_P5_C5
Global ipTM = 0.87
HC/Hu-1 RBD ipTM = 0.84
LC/Hu-1 RBD ipTM = 0.85
Global ipTM = 0.88
HC/Hu-1 RBD ipTM = 0.87
LC/Hu-1 RBD ipTM = 0.87
Global ipTM = 0.89
HC/Hu-1 RBD ipTM = 0.88
LC/Hu-1 RBD ipTM = 0.87
pLDDT confidence score
100 90 70 <50
b
All Hu-1 RBD and Fab/Fv predictions (n = 133)
HC ipTM
LC ipTM
Fv ipTM confidence score
Fab ipTM confidence score
Selected predictions (n=62)
c
All predictions
### of predictions
% of predictions (Fv)
% of predictions (Fab)
n = 133 n = 79 n = 54 n = 22 n = 12 n = 9 n = 8 n = 7 n = 6 n = 5 n = 4 n = 3 n = 2 n = 1
All sequences
Beta^high
Beta^low
IGHV1-69 IGHV3-30 IGHV1-58 IGHV3-66 IGHV3-9 IGHV2-5 IGHV1-3 IGHV2-23 IGHV5-51 IGHV1-18 IGHV1-46 IGHV3-30-3 IGHV3-33 IGHV1-2 IGHV3-53 IGHV3-15 IGHV3-49 IGHV3-64 IGHV3-7 IGHV4-39 IGHV4-61 IGHV3-11 IGHV3-13 IGHV4-31 IGHV5-10-1
Very high confidence (HC & LC ipTMs ≥0.8)
Confident (HC & LC ipTMs ≥ 0.6)
Poor confidence
d
Included in epitope binning
Fv
Fab
n = 86 n = 8 n = 15 n = 29 n = 8 n = 5 n = 21
% of predictions
All sequences A (1a) B (1b) D (2/3) E (3) C (4) F (4/5)
Epitope-binning group (corresponding class)
e
AF3 class
MBC group
Hu-1 RBD aligment
V320 P330 E340 V350 N360 N370 Y380 L390 F400 I410 D420 T430 N440 N450 N460 T470 C480 F490 T500 V510 A520 S530 N540
Class 2/3
Class 3
Class 1a
Class 1b
Class 4
Class 5a
Class 5b
ADG2
REGN10933
EY6A
S2H97
ACE2 footprint
RBD mutations
B.1.351
BA.1
G339D S371L S373P S375F K417N N440K G446S S477N T478K E484K/A Q493R G496S Q498R N501Y Y505H
MBC group
Beta^high
Beta^low
Reference mAbs
AlphaFold contact probability
1 0.5 0

**Fig. S4 (relates to Fig. 3). AlphaFold 3 predictions quality measurements for all anti-Hu-1 RBD $\mathbf{Beta^{high}}$ and $\mathbf{Beta^{low}}$ mAbs.**

(**a**) Representative Fv-Hu-1 RBD interaction model for each class as predicted by AlphaFold 3, with mAb identification (id), global (all three chains) and local ipTMs (heavy (HC) or light chain (LC) with the Hu-1 RBD) shown above and below. (**b**) Heavy (top) or light (bottom) chains local ipTM scores in interaction with the Hu-1 RBD for each tested mAb in their Fv versus Fab (Fv + IgG1 $C_{H1}$ domain) conformation. mAbs selected for downstream analysis, both HC and LC local ipTMs $\geq$ 0.6, are colored green. Fv models gave consistently higher iPTM scores and were used preferentially for epitope prediction analysis, except for 6, where Fab modeling was superior. Only Fab-based predictions were used for accessibility and bivalency predictions. (**c** and **d**) Percentage of very high confidence (white, both local ipTMs $\geq$ 0.8), confident (grey, both local ipTMs $\geq$ 0.6) or poor confidence predictions (black, one or more local ipTM $\leq$ 0.6) for Fv and Fab-based prediction of all mAbs in our dataset grouped according to $IgV_H$ gene usage (**c**) or the experimentally defined epitope-binning group (**Fig. 3a**) (**d**). Numbers of predictions for each group is further indicated on top of each panel. (**e**) Heatmap of predicted contact probabilities with individual Hu-1 RBD residues for all IgH/IgL pairs in our dataset and three published anti-RBD reference mAbs (class 1/4 ADG2, class 1 REGN10933, class 4 EY6A and class 5 S2H97) with "confident" to "very high confidence" AlphaFold 3 predictions. Unsupervised hierarchical clustering grouped mAbs into seven distinct antibody classes. The MBC population of origin for each mAb ($Beta^{low}$ or $Beta^{high}$) is indicated on the left side of the heatmap. The ACE2 footprint on the RBD and the RBD mutations in B.1.351 and/or BA.1 SARS-CoV-2 are shown below the heatmap.

Fig. S5

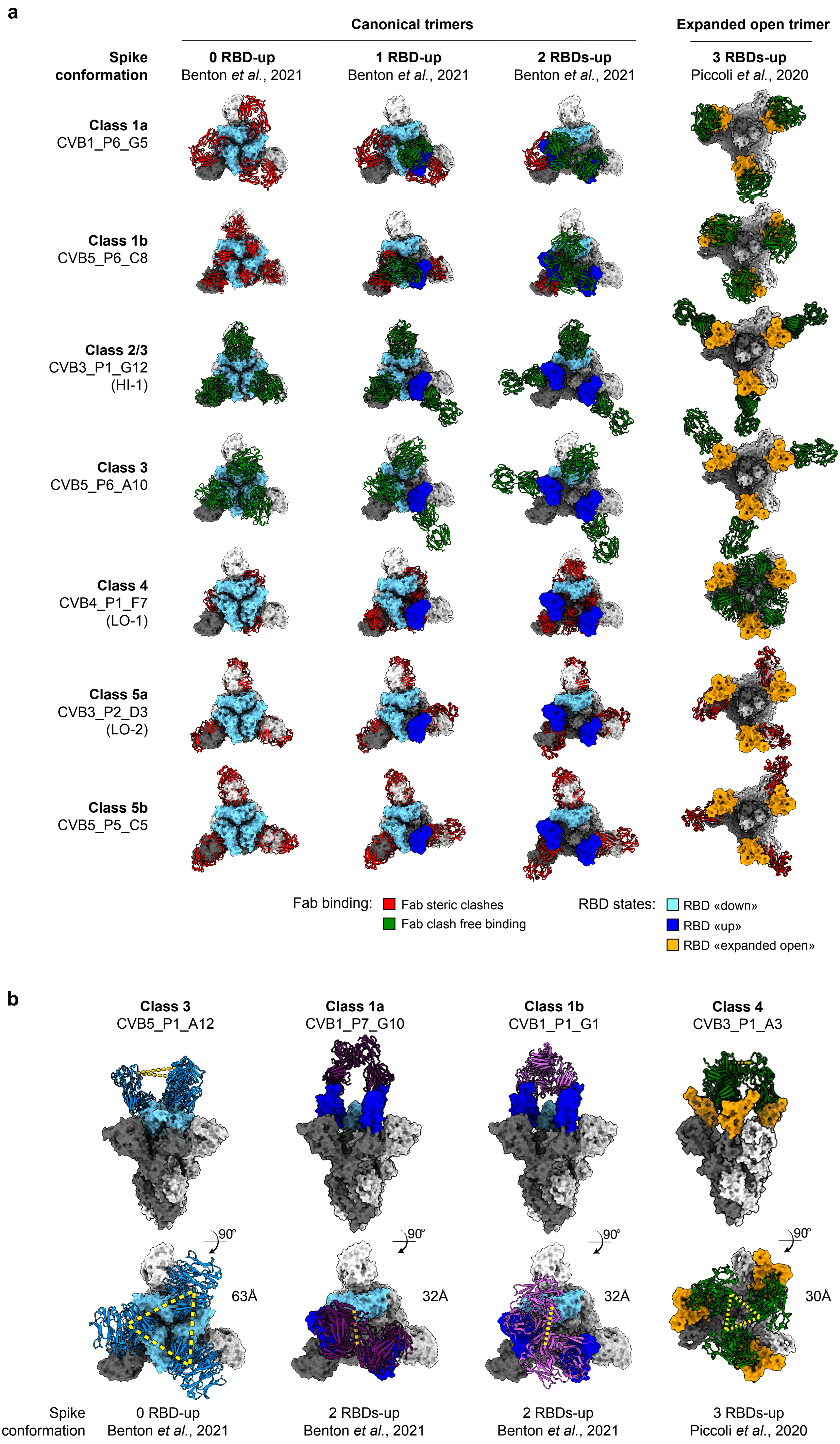
a
Canonical trimers
Expanded open trimer
Spike conformation
0 RBD-up
Benton et al., 2021
1 RBD-up
Benton et al., 2021
2 RBDs-up
Benton et al., 2021
3 RBDs-up
Piccoli et al., 2020
Class 1a
CVB1_P6_G5
Class 1b
CVB5_P6_C8
Class 2/3
CVB3_P1_G12
(HI-1)
Class 3
CVB5_P6_A10
Class 4
CVB4_P1_F7
(LO-1)
Class 5a
CVB3_P2_D3
(LO-2)
Class 5b
CVB5_P5_C5
Fab binding:
Fab steric clashes
Fab clash free binding
RBD states:
RBD «down»
RBD «up»
RBD «expanded open»
b
Class 3
CVB5_P1_A12
Class 1a
CVB1_P7_G10
Class 1b
CVB1_P1_G1
Class 4
CVB3_P1_A3
90°
63Å
32Å
32Å
30Å
Spike conformation
0 RBD-up
Benton et al., 2021
2 RBDs-up
Benton et al., 2021
2 RBDs-up
Benton et al., 2021
3 RBDs-up
Piccoli et al., 2020

**Fig. S5 (relates to Fig. 4). Structural modeling of RBD epitopes accessibility and mAbs bivalent binding potential in multiple prefusion D614G SARS-CoV-2 spike trimer conformational states.** (**a**) Alignments of Fab-RBD AlphaFold 3-predicted models for 7 representative mAbs from our datasets (one per identified mAb class) onto each monomer of indicated published cryo-EM spike trimer conformational state structure. Alignments were performed onto three canonical conformational states of the D614G SARS-CoV-2 spike protein, with zero (PDB 7BNM), one (PDB 7BNN) or two (PDB 7BNO) RBDs in the "up" state, alongside with an expanded open trimer stabilized upon antibody binding (S304, PDB 7JW0). RBD monomers are colored in light blue, dark blue and yellow when in the "down", "up" or "expanded open" position, respectively. Fabs are colored in red when steric clashes could be detected or in green in the case of clash-free binding. (**b**) Representative bivalency measurements for the 4 mAb class where bivalent binding could be predicted. Bivalency was assessed *in silico* across the four D614G SARS-CoV-2 spike trimer conformations described in (a) and bivalent binding was considered possible when the distance between the last $C_{H1}$ lysines of two bound Fabs to the same trimer was ≤ 65 Å (Barnes et al., 2020).

**Fig. S6**

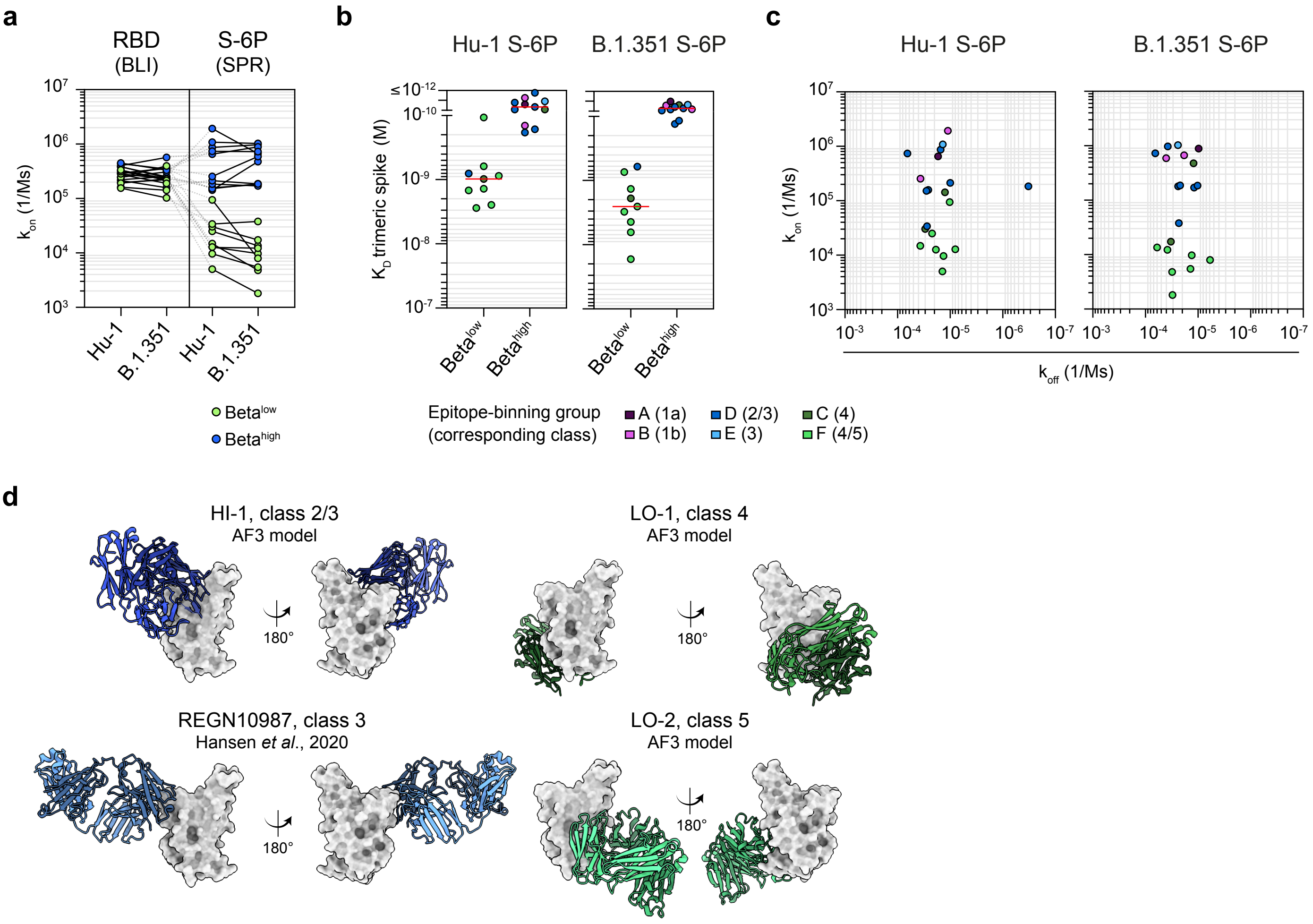

a
RBD (BLI)
S-6P (SPR)
k_on (1/Ms)
Hu-1
B.1.351
Beta^low
Beta^high
b
Hu-1 S-6P
B.1.351 S-6P
K_D trimeric spike (M)
c
k_on (1/Ms)
k_off (1/Ms)
Epitope-binning group (corresponding class)
A (1a)
B (1b)
D (2/3)
E (3)
C (4)
F (4/5)
d
HI-1, class 2/3
AF3 model
LO-1, class 4
AF3 model
REGN10987, class 3
Hansen et al., 2020
LO-2, class 5
AF3 model
180°

**Fig. S6 (relates to Figure 5). Extended analyses of affinity measurements onto the D614G and B.1.351 prefusion spike trimer (S-6P).** (**a**) Association rate constants $k_{on}$ (1/Ms) against the Hu-1 and B.1.351 RBD (measured by BLI) and Hu-1 and B.1.351 6-proline stabilized prefusion spike trimer (S-6P) (measured by Surface Plasmon Resonance (SPR)) for all $Beta^{high}$ (blue) and $Beta^{low}$ (green) tested mAbs. (**b-c**) Equilibrium dissociation constants ($K_D = k_{on}/k_{off}$, in M) measured by SPR (**c**) and scatter plot of association rate constants ($k_{on,}$, 1/Ms) vs dissociation rate constants ($k_{off}$, 1/s) against the full trimeric Hu-1 (left) or B.1.351 S-6P spike (right) for tested $Beta^{low}$ (n = 9) and $Beta^{high}$ (n = 11) mAbs. Tested mAbs were randomly selected among mAbs displaying similar affinity for Hu-1 and B.1.351 RBD and are further colored based on their experimentally determined epitope binning group (**Fig. 3a**). (**d**) AlphaFold 3-predicted Fab-Hu-1 RBD interaction models for the three re-expressed $Beta^{high}$ MBC-derived (class 2/3 CVB3_P1_G12, hereafter referred to as HI-1) and $Beta^{low}$ MBC-derived mAbs (class 4 CVB4_P1_F7 and class 5a CVB3_P2_D3, hereafter referred to as LO-1 and LO-2 respectively). The resolved structure of REGN10987 (PDB 6XDG) is also represented.
